# Sensitivity analysis of Wnt *β*-catenin based transcription complex might bolster power-logarithmic psychophysical law & reveal preserved gene gene interactions

**DOI:** 10.1101/015834

**Authors:** Shriprakash Sinha

**Author notes:** Code with dataset is made available under GNU GPL v3 license at google code project on https://sites.google.com/site/shriprakashsinha/shriprakashsinha/projects/static-bn-for-wnt-signaling-pathway. Please use the scripts in R as well as the files in zipped directory titled Results-2015. Email address (shriprakash sinha). This work was conducted by Sinha as an independent researcher during the period from 2013 to 2016. ORCID : orcid.org/0000-0001-7027-5788.

## Abstract

Recently, psychophysical laws have been observed to be functional in certain factors working downstream of the Wnt pathway. This work tests the veracity of the prevalence of such laws, albeit at a coarse level, using sensitivity analysis on biologically inspired epigenetically influenced computational causal models. In this work, the variation in the effect of the predictive behaviour of the transcription complex (TRCMPLX) conditional on the evidences of gene expressions in normal/tumor samples is observed by varying the initially assigned values of conditional probability tables (cpt) for TRCMPLX. Preliminary analysis shows that the variation in predictive behaviour of TRCMPLX follows power-logarithmic psychophysical law, crudely. More recently, wet lab experiments have proved the existence of sensors that behave in a logarithmic fashion thus supporting the earlier proposed postulates based on computational sensitivity analysis of this manuscript regarding the existence of logarithmic behaviour in the signaling pathways. It also signifies the importance of systems biology approach where in silico experiments combined with in vivo/in vitro experiments have the power to explore the deeper mechanisms of a signaling pathway. Additionally, it is hypothesized that these laws are prevalent at gene-gene interaction level also. The interactions were obtained by thresholding the inferred conditional probabilities of a gene activation given the status of another gene activation. The deviation in the interactions in normal/tumor samples was similarly observed by varying the initially assigned values of conditional probability tables (cpt) for TRCMPLX. Analysis of deviation in interactions show prevalence of psychophysical laws and is reported for interaction between elements of pairs (SFRP3, MYC), (SFRP2, CD44) and (DKK1, DACT2). Based on crude static models, it is assumed that dynamic models of Bayesian networks might reveal the phenomena in a better way.

## 1. Introduction & problem statement

Ever since the accidental discovery of the Wingless in 1973 by Sharma (1973), a tremendous amount of research work has been carried out in the related field of Wnt signaling pathway in the past forty years. A majority of the work has focused on issues related to • the discovery of genetic and epigenetic factors affecting the pathway Thorstensen et al. (2005) & Baron and Kneissel (2013), • implications of mutations in the pathway and its dominant role on cancer and other diseases Clevers (2006), • investigation into the pathway’s contribution towards embryo development Sokol (2011), homeostasis Pinto et al. (2003), Zhong et al. (2014) and apoptosis Pećina-Šlaus (2010) and • safety and feasibility of drug design for the Wnt pathway Kahn (2014), Garber (2009), Voronkov and Krauss (2012), Blagodatski et al. (2014) & Curtin and Lorenzi (2010). More recent informative reviews have touched on various issues related to the different types of the Wnt signaling pathway and have stressed not only the activation of the Wnt signaling pathway via the Wnt proteins Rao and Kühl (2010) but also the on the secretion mechanism that plays a major role in the initiation of the Wnt activity as a prelude Yu and Virshup (2014).

In a more recent development, there has been the observation and study of psychophysical laws prevailing within the pathway and in this regard Goentoro and Kirschner (2009) point to two findings namely, • the robust fold changes of *β*-*catenin* • and the transcriptional machinery of the Wnt pathway depends on the fold changes in *β*-*catenin* instead of absolute levels of the same and some gene transcription networks must respond to fold changes in signals according to the Weber (1834) law in sensory physiology. Note that Weber’s law has been found to be a special case of Bernoulli’s logarithmic law Masin et al. (2009). If a sensation magnitude *γ* be determined by a stimulus magnitude *β*, then the Weber’s law states that Δ*γ* remains constant when the relative stimulus increment Δ*β* remains constant. The law derives from a more general Bernoullis law were Δ*γ* ∝ log 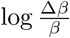. In an unrelated work by Sun et al. (2012), it has been shown that these laws arise at computational level as Bayes optimal under neurobiological constraints at implementational and algorithmic levels. The proposed mathematical framework for understanding the psychophysical scales as Bayes optimal and information theoretically-optimal representation of time sampled continuous valued stimuli is based on established neurobiological assumptions. Sun et al. (2012) also show that the psychophysical laws connect well to quantization frameworks and state that only discrete set of output is distinguishable due to biological constraints. This discretization leads to quantization of stimulus also as the nonlinear scaling of the stimulus that leads to the resultant output is invertible. These mathematical insights might explain the indistinguishable insensitive fold changes in levels of *β*-*catenin* shown by Goentoro and Kirschner (2009).

Based on the importance of the revealed phenomena, it might be useful to know if these observations could be verified using computational models apart from analysis of results from wet lab experiments. What is needed is a frame-work that can capture the causal semantics of the signaling pathway where the influence diagrams involving the interacting extra/intracellular factors working in the pathway, represent the biological knowledge/mechanism of the pathway to a certain extent. Once a model representation is available, the desired variation in the activity of an input factor and the observed variation in the output of the activity of factor(s) can be studied. Sensitivity analysis plays a crucial role in observing the behaviour of output of a variable given variations in the input. As will be seen later, probabilistic graphical models or Bayesian networks provide a framework for representing the causal semantics of the pathway under investigation.

To address these issues, the current work uses the Bayesian network model proposed in Sinha (2014) and conducts sensitivity analysis on the model to check the observations regarding the prevalence of the reported psychophysical laws. In Sinha (2014), it was shown via hypothesis testing that the active (repressed) state of *TRCMPLX* in the Wnt signaling pathway for colorectal cancer cases is not always correspond to the tumorous (normal) state of the test sample under consideration. For this, Sinha (2014) shows various results on the predicted state of *TRCMPLX* conditional on the given gene evidences, while varying the assigned probability values of conditional probability tables of *TRCMPLX* during initialization of the biologically inspired Bayesian Network model. Here, the degree of belief in the activity of *TRCMPLX* is denoted by the prior probability assigned to the node of *TRCMPLX* in the network. It was found that the predicted values often increase with an increasing value (in conditional probability tables) of the activity of *TRCMPLX* on certain genes. What this asks for is that for the recorded deviations due to the changes made in these prior probabilities (i.e the input deviations), is it possible to observe the prevalent logarithmic laws and their deviations (like the Weber’s law) as shown by Goentoro and Kirschner (2009), using computational causal modeling?

In this manuscript, the preliminary analysis of deviations computed from variation in prior and estimated conditional probability values using Bayesian network model in Sinha (2014) show that the variation in predictive behaviour of *TRCMPLX* conditional on gene evidences (i.e the output deviation) follows power and logarithmic psychophysical law crudely, apropos the variation in assigned priors of *TRCMPLX* (i.e input deviations). This implies that the deviations in output are proportional to increasing function of deviations in input. This relates to the work of Adler et al. (2014) on power and logarithmic law albeit at a coarse level. The granularity is obscured due to the use of static data from Jiang et al. (2008) that is used in Sinha (2014) as well as the Bayesian network model that encodes the belief in the factors affecting the pathway in terms of probabilities as well as the inferences made based on the updating of these probabilities conditional on discretized states of gene expression values as evidences. Irrespective of the hurdle posed by the causal models, inferences made based on prior biological knowledge and gene expression evidences coupled with sensitivity analysis sheds light on the prevalent power-logarithmic psychophysical laws in the pathway. *Note that the foundations of the current work were presented as poster in the International Conference on Systems Biology of Human Disease at the German Cancer Research Center in Heidelberg (Germany) in 2015. Followup of some of the implications were shared with a few labs for verification and it is gladdening to see that in a recent development via wet lab experiments by Olsman and Goentoro (2016), it has been confirmed that there are existence of sensors that behave in a logarithmic fashion. The wet lab work by Olsman and Goentoro (2016) supports the earlier proposed crude postulates based on computational sensitivity analysis of this manuscript regarding the existence of logarithmic behaviour in the signaling pathways. It also signifies the importance of systems biology approach where in silico experiments combined with in vivo/in vitro experiments have the power to explore the deeper mechanisms of a signaling pathway*.

Adler et al. (2014) show in detail that these laws can be studied empirically using models that exhibit the property of fold change detection (FCD). What this means is that the output depends on the relative changes in the input. The biological feedback models employed for these studies consider various parameters like rates of production of a compound, removal removal of a compound, repression of a compound, levels of scaffolds, kinases, etc. that might be responsible for exhibiting these laws. The current work using the static Bayesian network model might not propose feedback loops directly as used by Adler et al. (2014), yet it could reveal existence of the loops via causal inference even while using static data. The drawback of the current work is its inability to consider cyclic loops. This can be rectified by use of dynamic Bayesian network models that incorporate interaction represented in time series data. Also, the use of Bayesian network models can help in studying the problem from a multiparameter setting as various factors affecting the pathway can be connected in the influence diagrams of the network through the principle of *d*-connectivity/separability. This connectivity will be explained later in the required theory section.

Note that Goentoro and Kirschner (2009) show results for the behaviour of fold change of *β*-*catenin* with respect to changes in the single parameter values i.e the Wnt. On similar lines, the current work takes into account the behaviour of *TRCMPLX* conditional on affects of multiple parameters in the form of evidences of various intra/extracellular gene expression values working in the pathway, based on the changes made in the assigned prior probabilities for *TRCMPLX*. The difference here is that one can analyse changes in nodes of a computational model to explore an inherent law in comparison to use of wet lab experiments. The issue here is that FCD which is recorded with respect to changes in levels of concentration can now be recorded via changes in the strength of belief in the occurrence of an event. For example, suppose it is not known by what degree the *TRCMPLX* plays a major role in the signaling pathway quantitatively, then it is possible to encode the degree of belief regarding the role of *TRCMPLX* in the form of prior or conditional probabilities during initialization of the network. By recording the deviations in these probabilities and observing the output deviations, it is possible to study certain psychophysical laws. Finally, this does not mean that probabilities related to actual concentrations cannot be encoded. Thus, Bayesian networks help in capturing the desired biological knowledge via various causal arcs and conditional probabilities and sensitivity analysis aids in the study of the such natural behaviour.

As a second observation, the forgoing result points towards stability in the behaviour of *TRCMPLX* and this stability is reflected in the preserved gene gene interactions across the changing values of the priors of *TRCMPLX*. The interactions are inferred from conditional probabilities of individual gene activation given the status of another gene activation. Finally, as a third observation, it would be interesting to note if the psychophysical laws are prevalent among the dual gene-gene interactions or not. If the results are affirmative then the following important speculations might hold true • Not just one factor but components of the entire network might be exhibiting such a behavior at some stage or the other. • The psychophysical law might not be restricted to individual intra/extracellular components but also to the interactions among the the intra/extracellular components in the pathway. This might mean that the interactions manifest during the prevalence of power and logarithmic laws. Further wet lab analysis is needed to very these computational claims.

It is important to be aware of the fact that the presented results are derived from a static Bayesian network model. It is speculated that dynamic models might give much better and more realistic results.

## 2. Revisiting the requisite theory

To understand the logical flow of the current paper, some details of the above related topics from Sinha (2014) are revisited here in order and subdivided into descriptions of - (1) general working of canonical Wnt signaling pathway and some of the involved epigenetic factors (2) introduction to Bayesian networks and (3) the intuition behind the Bayesian network model employed. This is followed by Weber’s law and its derivation and finally the notations and terminologies to understand the results and discussion section.

### 2.1 Canonical Wnt signaling pathway

The canonical Wnt signaling pathway is a transduction mechanism that contributes to embryo development and controls homeostatic self renewal in several tissues Clevers (2006). Somatic mutations in the pathway are known to be associated with cancer in different parts of the human body. Prominent among them is the colorectal cancer case Gregorie (2005) and Clevers (2005). In a succinct overview, the Wnt signaling pathway works when the Wnt ligand gets attached to the Frizzled(*fzd*)/*LRP* coreceptor complex. *Fzd* may interact with the Dishevelled (*Dvl*) causing phosphorylation. It is also thought that Wnts cause phosphorylation of the *LRP* via casein kinase 1 (*CK*1) and kinase *GSK*3. These developments further lead to attraction of Axin which causes inhibition of the formation of the degradation complex. The degradation complex constitutes of *Axin*, the *β*-*catenin* transportation complex *APC*, *CK*1 and *GSK*3. When the pathway is active the dissolution of the degradation complex leads to stabilization in the concentration of *β*-*catenin* in the cytoplasm. As *β*-*catenin* enters into the nucleus it displaces the *Groucho* and binds with transcription cell factor *TCF* thus instigating transcription of Wnt target genes. *Groucho* acts as lock on *TCF* and prevents the transcription of target genes which may induce cancer. In cases when the Wnt ligands are not captured by the coreceptor at the cell membrane, *Axin* helps in formation of the degradation complex. The degradation complex phosphorylates *β*-*catenin* which is then recognized by *Fbox*/*WD* repeat protein *β* − *TrCP*. *β* − *TrCP* is a component of ubiquitin ligase complex that helps in ubiquitination of *β*-*catenin* thus marking it for degradation via the proteasome. Cartoons depicting the phenomena of Wnt being inactive and active are shown in figures 1(A) and 1(B), respectively.

**Figure 1:**
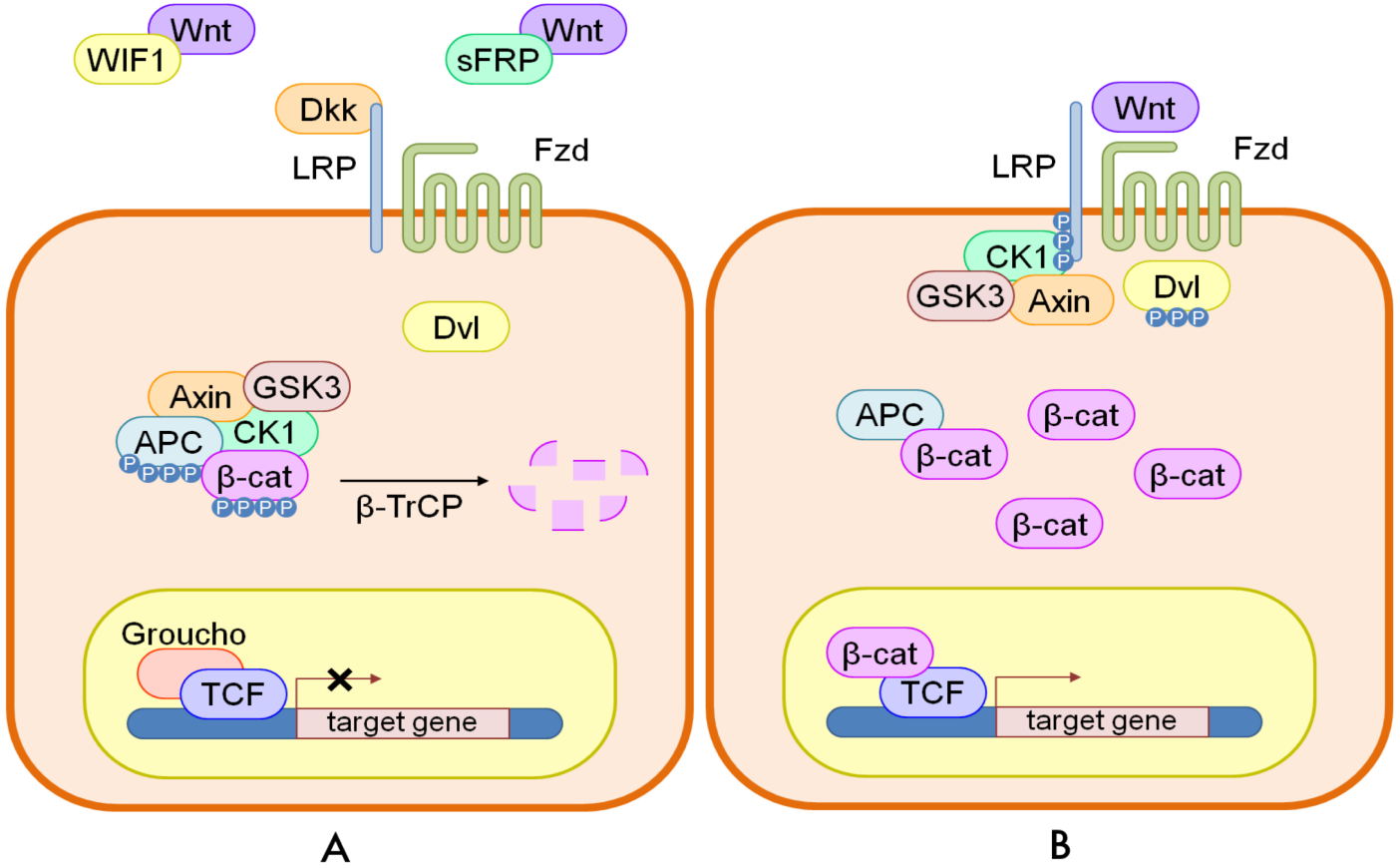
A cartoon of Wnt signaling pathway adapted from Sinha (2014). Part (A) represents the destruction of *β*-*catenin* leading to the inactivation of the Wnt target gene. Part (B) represents activation of Wnt target gene.

### 2.2 Epigenetic factors

One of the widely studied epigenetic factors is methylation Costello and Plass (2001), Das and Singal (2004), Issa (2007). Its occurrence leads to decrease in the gene expression which affects the working of Wnt signaling pathways. Such characteristic trends of gene silencing like that of secreted frizzled-related proteins (*SFRP*) family in nearly all human colorectal tumor samples have been found at extracellular level Suzuki et al. (2004). Similarly, methylation of genes in the Dickkopf (*DKKx* Niehrs (2006), Sato et al. (2007), Dapper antagonist of catenin (*DACTx* Jiang et al. (2008) and Wnt inhibitory factor-1 (*WIF* 1 Taniguchi et al. (2005) family are known to have significant effect on the Wnt pathway. Also, histone modifications (a class of proteins that help in the formation of chromatin which packs the DNA in a special form Strahl and Allis (2000) can affect gene expression Peterson et al. (2004). In the context of the Wnt signaling pathway it has been found that *DACT* gene family show a peculiar behavior in colorectal cancer Jiang et al. (2008). *DACT* 1 and *DACT* 2 showed repression in tumor samples due to increased methylation while *DACT* 3 did not show obvious changes to the interventions. It is indicated that *DACT* 3 promoter is simultaneously modified by the both repressive and activating (bivalent) histone modifications Jiang et al. (2008).

### 2.3 Bayesian Networks

In reverse engineering methods for control networks Gardner and Faith (2005) there exist many methods that help in the construction of the networks from the data sets as well as give the ability to infer causal relations between components of the system. A widely known architecture among these methods is the Bayesian Network (BN). These networks can be used for causal reasoning or diagnostic reasoning or both. It has been shown through reasoning and examples in Roehrig (1996) that the probabilistic inference mechanism applied via Bayesian networks are analogous to the structural equation modeling in path analysis problems.

Initial works on BNs in Pearl (1988) and Pearl (2000) suggest that the networks only need a relatively small amount of marginal probabilities for nodes that have no incoming arcs and a set of conditional probabilities for each node having one or more incoming arcs. The nodes form the driving components of a network and the arcs define the interactive influences that drive a particular process. Under these assumptions of influences the joint probability distribution of the whole network or a part of it can be obtained via a special factorization that uses the concept of direct influence and through dependence rules that define d-connectivity/separability as mentioned in Charniak (1991) and Needham et al. (2007). This is illustrated through a simple example in Roehrig (1996).

The Bayesian networks work by estimating the posterior probability of the model given the data set. This estimation is usually referred to as the Bayesian score of the model conditioned on the data set. Mathematically, let 𝒮 represent the model given the data 𝒟 and *ξ* is the background knowledge. Then according to the Bayes Theorem Bayes and Price (1763):

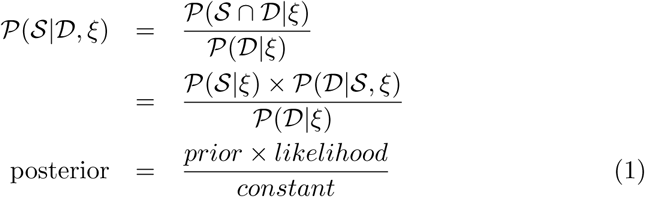

Thus the Bayesian score is computed by evaluating the **posterior** distribution *𝒫* (*𝒮| 𝒟, ξ*) which is proportional to the **prior** distribution of the model *𝒫* (*𝒮|ξ*) and the **likelihood** of the data given the model *𝒫* (*𝒟|* (*𝒮, ξ*). It must be noted that the background knowledge is assumed to be independent of the data. Next, since the evaluation of probabilities require multiplications a simpler way is to take logarithmic scores which boils down to addition. Thus the estimation takes the form:

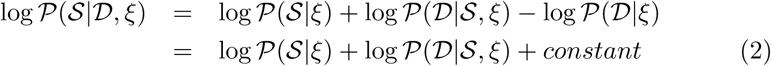

Finally, the likelihood of the function can be evaluated by averaging over all possible local conditional distributions parameterized by *θ*_*i*_’s that depict the conditioning of parents. This is equated via:

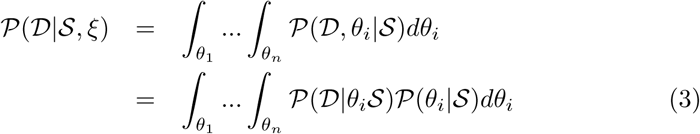

Work on biological systems that make use of Bayesian networks can also be found in Friedman et al. (2000), Hartemink et al. (2001), Sachs et al. (2002), Sachs et al. (2005) and Peer et al. (2001). Bayesian networks are good in generating network structures and testing a targeted hypothesis which confine the experimenter to derive causal inferences Brent and Lok (2005). But a major disadvantage of the Bayesian networks is that they rely heavily on the conditional probability distributions which require good sampling of datasets and are computationally intensive. On the other hand, these networks are quite robust to the existence of the unobserved variables and accommodates noisy datasets. They also have the ability to combine heterogeneous data sets that incorporate different modalities.

In Sinha (2014), simple static Bayesian Network models have been developed with an aim to show how • incorporation of heterogeneous data can be done to increase prediction accuracy of test samples • prior biological knowledge can be embedded to model biological phenomena behind the Wnt pathway in colorectal cancer *•* to test the hypothesis regarding direct correspondence of active state of *β*-*catenin* based transcription complex and the state of the test sample via segregation of nodes in the directed acyclic graphs of the proposed models and • inferences can be made regarding the hidden biological relationships between a particular gene and the *β*-*catenin* transcription complex. This work uses MATLAB implemented BN toolbox from Murphy et al. (2001).

### 2.4 Intuition behind the causal semantics of the biologically inspired Bayesian network

The NB model Ver assumes that the activation (inactivation) of *β*-*catenin* based transcription complex is equivalent to the fact that the sample is cancerous (normal). This assumption needs to be tested and Sinha (2014) proposes a newly improvised models based on prior biological knowledge and epigenetic information regarding the signaling pathway with the assumption that sample prediction may not always mean that the *β*-*catenin* based transcription complex is activated. These assumptions are incorporated by inserting another node of *Sample* for which gene expression measurements were available. This is separate from the *TRCMPLX* node that influences a particular set of known genes in the human colorectal cancer. For those genes whose relation with the *TRCMPLX* is currently not known or biologically affirmed, indirect paths through the *Sample* node to the *TRCMPLX* exist, technical aspect of which is described next.

For those factors whose relations were not yet confirmed but known to be involved in the pathway, the causal arcs were segregated via a latent variable introduced into the Bayesian network. The latent variable in the form of *Sample* (see figure 2) is extremely valuable as it connects the factors whose relations have not been confirmed till now, to factors whose influences have been confirmed in the pathway. Finally, the introduction of latent variable in a causal model opens the avenue to assume the presence of measurements that haven’t been recorded. Intuitively, for cancer samples the hidden measurements might be different from those for normal samples. The connectivity of factors through the variable provides an important route to infer biological relations.

**Figure 2:**
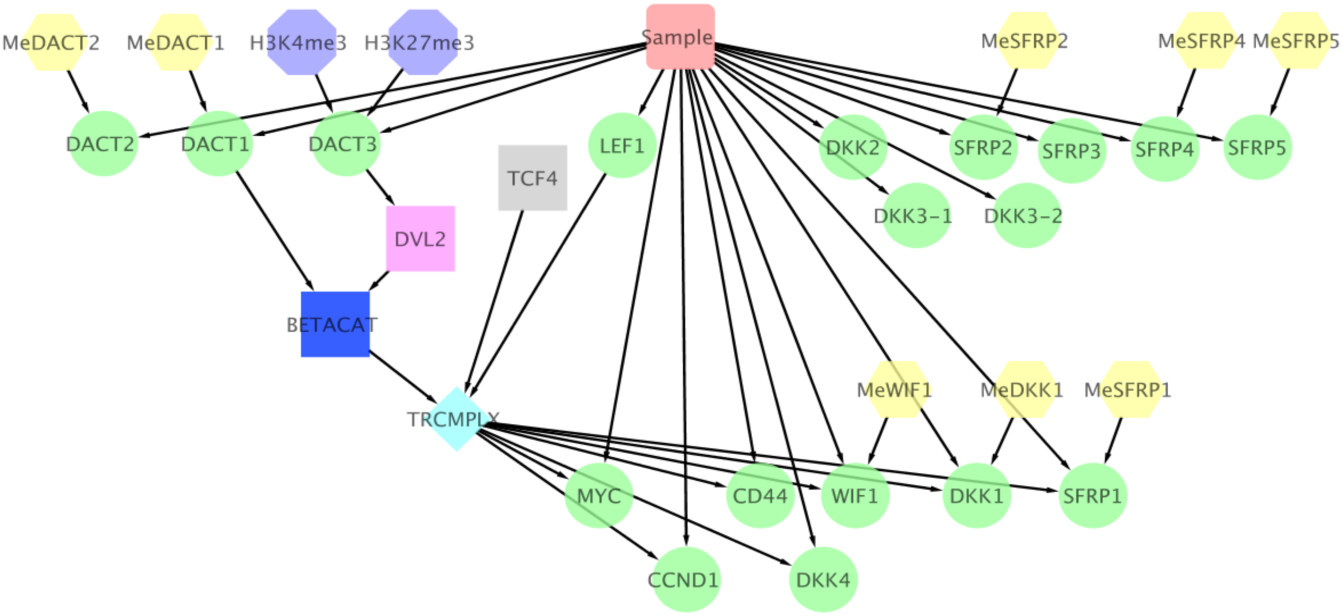
Influence diagram of *M*_*PBK*+*EI*_ contains partial prior biological knowledge and epigenetic information in the form of methylation and histone modification. Diagram drawn using CYTOSCAPE Shannon et al. (2003). In this model the state of Sample is distinguished from state of *TRCMPLX* that constitutes the Wnt pathway.

Since all gene expressions have been measured from a sample of subjects the expression of genes is conditional on the state of the *Sample*. Here both tumorous and normal cases are present in equal amounts. The transcription factor *TRCMPLX* under investigation is known to operate with the help of interaction between *β*-*catenin* with *TCF* 4 and *LEF* 1 Waterman (2004), Kriegl et al. (2010). It is also known that the regions in the TSS of *MY C* Yochum *(2011), CCND*1 Schmidt-Ott et al. (2007), *CD*44 Kanwar et al. (2010), *SFRP* 1 Caldwell et al. (2006), *WIF* 1 Reguart et al. (2004), *DKK*1 González-Sancho et al. (2004) and *DKK*4 Pendas-Franco et al. (2008), Baehs et al. (2009) contain factors that have affinity to *β*-*catenin* based *TRCMPLX*. Thus expression of these genes are shown to be influenced by *TRCMPLX*, in figure 2.

Roles of *DKK*2 Matsui et al. (2009) and *DKK*3 Zitt et al. (2008), Veeck and Dahl (2012) have been observed in colorectal cancer but their transcriptional relation with *β*-*catenin* based *TRCMPLX* is not known. Similarly, *SFRP* 2 is known to be a target of *Pax*2 transcription factor and yet it affects the *β*-*catenin* Wnt signaling pathway Brophy et al. (2003). Similarly, *SFRP* 4 Feng Han et al. (2006), Huang et al. (2010) and *SFRP* 5 Suzuki et al. (2004) are known to have affect on the Wnt pathway but their role with *TRCMPLX* is not well studied. *SFRP* 3 is known to have a different structure and function with respect to the remaining *SFRPx* gene family Hoang et al. (1996). Also, the role of *DACT* 2 is found to be conflicting in the Wnt pathway Kivimäe et al. (2011). Thus for all these genes whose expression mostly have an extracellular affect on the pathway and information regarding their influence on *β*-*catenin* based *TRCMPLX* node is not available, an indirect connection has been made through the *Sample* node. This connection will be explained at the end of this section.

Lastly, it is known that concentration of *DV L*2 (a member of Disheveled family) is inversely regulated by the expression of *DACT* 3 Jiang et al. (2008). High *DV L*2 concentration and suppression of *DACT* 1 leads to increase in stabilization of *β*-*catenin* which is necessary for the Wnt pathway to be active Jiang et al. (2008). But in a recent development Yuan et al. (2012) it has been found that expression of *DACT* 1 positively regulates *β*-*catenin*. Both scenarios need to be checked via inspection of the estimated probability values for *β*-*catenin* using the test data. Thus there exists direct causal relations between parent nodes *DACT* 1 and *DV L*2 and child node, *β*-*catenin*. Influence of methylation (yellow hexagonal) nodes to their respective gene (green circular) nodes represent the affect of methylation on genes. Influence of histone modifications in *H*3*K*27*me*3 and *H*3*K*4*me*3 (blue octagonal) nodes to *DACT* 3 gene node represents the affect of histone modification on *DACT* 3. The *β*-*catenin* (blue square) node is influenced by concentration of *DV L*2 (depending on the expression state of *DACT* 3) and behavior of *DACT* 1.

The aforementioned established prior causal biological knowledge is imposed in the Bayesian network model with the aim to computationally reveal unknown biological relationships. The influence diagram of this model is shown in figure 2 with nodes on methylation and histone modification.

In order to understand indirect connections further it is imperative to know about **d-connectivity/separability**. In a BN model this connection is established via the principle of **d-connectivity** which states that nodes are *connected* in a path when there exists no node in the path that has more than one incoming influence edge or there exits nodes in path with more than one incoming influence edge which are observed (i.e evidence regarding such nodes is available) Charniak (1991). Conversely, via principle of **d-separation** nodes are *separated* in a path when there exists nodes in the path that have more than one incoming influence edge or there exists nodes in the path with at most one incoming influence edge which are observed (i.e evidence regarding such nodes is available). Figure 3 represents three different cases of connectivity and separation between nodes 𝒜 and 𝒞 when the path between them passes through node. ℬ Connectivity or dependency exists between nodes 𝒜 and 𝒞 when • evidence is not present regarding node ℬ in the left graphs of I. and II. in figure 3 or • evidence is present regarding node ℬ in the right graph of III. in figure Conversely, separation or independence exits between nodes 𝒜 and 𝒞 when • evidence is present regarding node ℬ in the right graphs of I. and II. in figure 3 or • evidence is not present regarding node ℬin the left graph of III. in figure 3.

**Figure 3:**
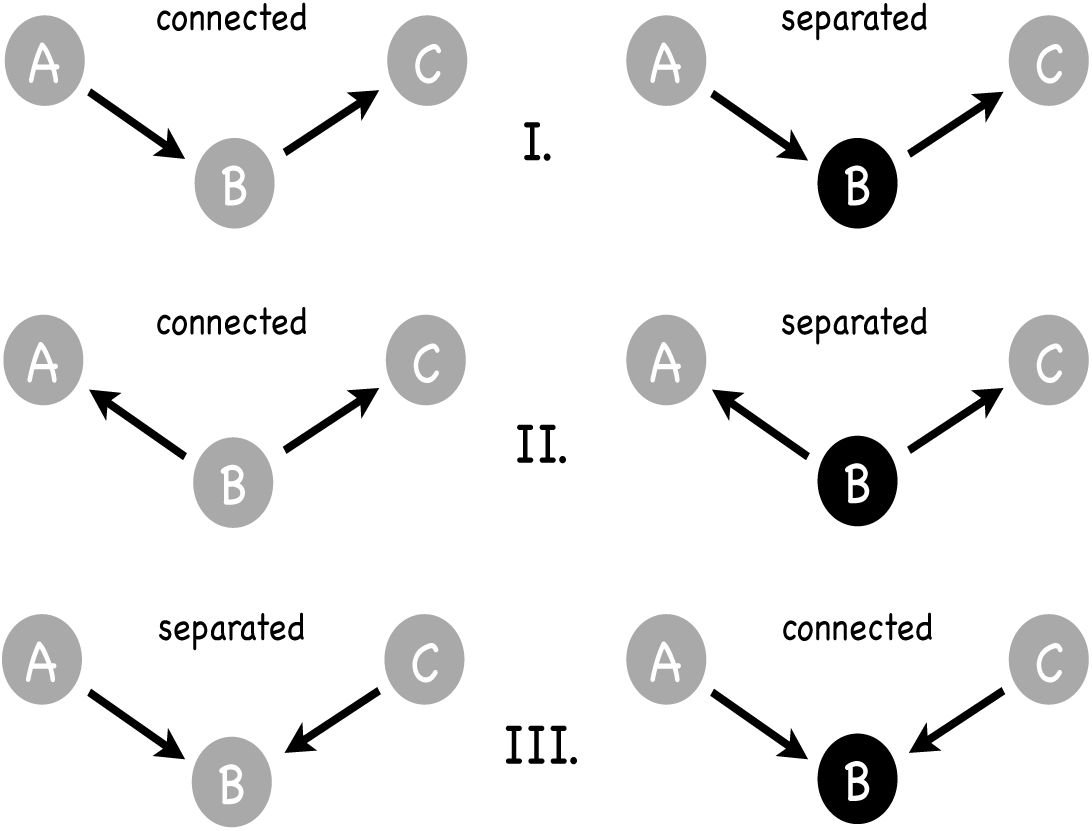
Cases for d-connectivity and d-separation. Black (Gray) circles mean evidence is available (not available) regarding a particular node.

It would be interesting to know about the behaviour of *TRCMPLX* given the evidence of state of *SFRP* 3. To reveal such information paths must exist between these nodes. It can be seen that there are multiple paths between *TRCMPLX* and *SFRP* 2 in the BN model in figure 2. These paths are enumerated as follows:

1. *SFRP* 3, *Sample*, *SFRP* 1, *TRCMPLX*
2. *SFRP* 3, *Sample*, *DKK*1, *TRCMPLX*
3. *SFRP* 3, *Sample*, *WIF* 1, *TRCMPLX*
4. *SFRP* 3, *Sample*, *CD*44, *TRCMPLX*
5. *SFRP* 3, *Sample*, *DKK*4, *TRCMPLX*
6. *SFRP* 3, *Sample*, *CCND*1, *TRCMPLX*
7. *SFRP* 3, *Sample*, *MY C*, *TRCMPLX*
8. *SFRP* 3, *Sample*, *LEF* 1, *TRCMPLX*
9. *SFRP* 3, *Sample*, *DACT* 3, *DV L*2, *β*-*catenin*, *TRCMPLX*
10. *SFRP* 3, *Sample*, *DACT* 1, *β*-*catenin*, *TRCMPLX*

Knowledge of evidence regarding nodes of *SFRP* 1 (path 1), *DKK*1 (path 2), *WIF* 1 (path 3), *CD*44 (path 4), *DKK*4 (path 5), *CCND*1 (path 6) and *MY C* (path 7) makes *Sample* and *TRCMPLX* dependent or d-connected. Further, no evidence regarding state of *Sample* on these paths instigates dependency or connectivity between *SFRP* 3 and *TRCMPLX*. On the contrary, evidence regarding *LEF* 1, *DACT* 3 and *DACT* 1 makes *Sample* (and child nodes influenced by *Sample*) independent or d-separated from *TRCMPLX* through paths (8) to (10). Due to the dependency in paths (1) to (7) and the given state of *SFRP* 3 (i.e evidence regarding it being active or passive), the BN uses these paths during inference to find how *TRCMPLX* might behave in normal and tumorous test cases. Thus, exploiting the properties of d-connectivity/separability, imposing a biological structure via simple yet important prior causal knowledge and incorporating epigenetic information, BN help in inferring many of the unknown relation of a certain gene expression and a transcription complex.

### 2.5. The logarithmic psychophysical law

Masin et al. (2009) states the Weber’s law as follows -

Consider a sensation magnitude *γ* determined by a stimulus magnitude *β*. Fechner (1860) (vol 2, p. 9) used the symbol Δ*γ* to denote a just noticeable sensation increment, from *γ* to *γ* + Δ*γ*, and the symbol Δ*β* to denote the corresponding stimulus increment, from *β* to *β* + Δ*β*. Fechner (1860) (vol 1, p. 65) attributed to the German physiologist Ernst Heinrich Weber the empirical finding Weber (1834) that Δ*γ* remains constant when the relative stimulus increment 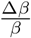 remains constant, and named this finding Weber’s law. Fechner (1860) (vol 2, p. 10) underlined that Weber’s law was empirical.

It has been found that Bernoulli’s principle Bernoulli (1738) is different from Weber’s law Weber (1834) in that it refers to Δ*γ* as any possible increment in *γ*, while the Weber’s law refers only to just noticeable increment in *γ*. Masin et al. (2009) shows that Weber’s law is a special case of Bernoulli’s principle and can be derived as follows - Equation 4 depicts the Bernoulli’s principle and increment in sensation represented by Δ*γ* is proportional to change in stimulus represented by Δ*β*.

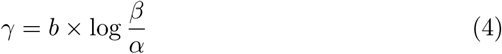

were *b* is a constant and *α* is a threshold. To evaluate the increment, the following Equation 5 and the ensuing simplification gives -

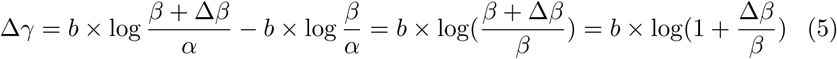

Since *b* is a constant, Equation 5 reduces to

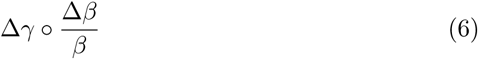

were *o* means ”is constant when there is constancy of” from Masin et al. (2009). The final Equation 6 is a formulation of Weber’s laws in wordings and thus Bernoulli’s principles imply Weber’s law as a special case. Using Fechner (1860) derivation, it is possible to show the relation between Bernoulli’s principles and Weber’s law. Starting from the last line of Equation 5, the following yields the relation.

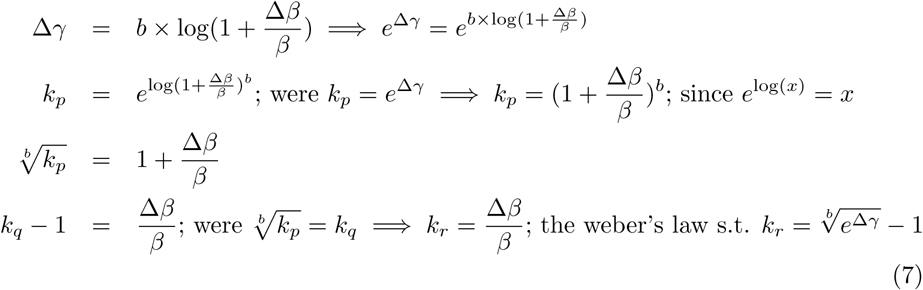

Equation 6 holds true given the last line of Equation 7. In the current study, observation of deviation recorded in predicted values of state of *TRCMPLX* conditional on gene evidences show crude logarithmic behaviour which might bolster Weber’s law and Bernoulli’s principles. But it must be noted that these observations are made on static causal models and observation of the same behaviour in dynamical setting would add more value.

## 3. Materials and methods

The models purported by Sinha (2014) involving the biological knowledge as well as epigenetic information depicted by *M*_*PBK*+*EI*_ and biological knowledge excluding epigenetic information *M*_*PBK*_ were used to predict the state of *TRCMPLX* given the gene evidences. Figure 2 depicts the model *M*_*PBK*+*EI*_. The predictions were recorded over the varying effect of *TRCMPLX* on gene regulations via assignment of different values to conditional probability tables (cpt) of *TRCMPLX* while initializing the aforementioned BN models. This varying effect is represented by the term ETGN in Sinha (2014).

As a recapitulation, the design of the experiment is a simple 2-holdout experiment where one sample from the normal and one sample from the tumorous are paired to form a test dataset. Excluding the pair formed in an iteration of 2-hold out experiment the remaining samples are considered for training of a BN model. Thus in a data set of 24 normal and 24 tumorous cases obtained from Jiang et al. (2008), a training set will contain 46 samples and a test set will contain 2 samples (one of normal and one of tumor). This procedure is repeated for every normal sample which is combined with each of the tumorous sample to form a series of test datasets. In total there will be 576 pairs of test data and 576 instances of training data. Note that for each test sample in a pair, the expression value for a gene is discretized using a threshold computed for that particular gene from the training set. Computation of the threshold has been elucidated in Sinha (2014). This computation is repeated for all genes per test sample. Based on the available evidence from the state of expression of all genes, which constitute the test data, inference regarding the state of both the *TRCMPLX* and the test sample is made. These inferences reveal information regarding the activation state of the *TRCMPLX* and the state of the test sample. Finally, for each gene *g*_*i*_, the conditional probability Pr(*g*_*i*_ = active | *g*_*k*_ evidence) *∀k* genes. Note that these probabilities are recorded for both normal and tumor test samples.

Three observations are presented in this manuscript. The **first observation** is regarding the logarithmic deviations in prediction of activation status of *TRCMPLX* conditional on gene expression evidences. The **second observation** is preservation of some gene gene interactions across different strength of beliefs concerning the affect of *TRCMPLX*. To observe these preservations, first the gene gene interactions have to be constructed from the predicted conditional probabilities of one gene given the evidence of another gene (for all gene evidences taken separately). After the construction, further preprocessing is required before the gene-gene interaction network can be inferred. Finally, the **third observation** is to check whether these laws are prevalent among the gene-gene interactions in the network or not.

## 4. Results and discussion on observation 1

### 4.1 Logarithmic-power deviations in predictions of β-catenin transcription complex

Let *γ* be Pr(*TRCMPLX* = active all gene evidences), *β* be the assigned cpt value of *TRCMPLX* during initialization of the Bayesian network models and Δ*β* be the deviation in the assigned values of *TRCMPLX* during initialization. To compute Δ*γ*, the 576 predictions of *γ* observed at *β* = 90% is subtracted from the 576 predictions of *γ* observed at *β* = 80% and a mean of the deviations recorded. This mean becomes Δ*γ*. The procedure is computed again for different value of *β*. In this manuscript, the effect of constant and incremental deviations are observed. Tables 1 and 2 represent the deviations for models *M* _*PBK*+*EI*_ and *M* _*PBK*_, respectively.

**Table 1:** Deviation study for model *M*_*PBK*+*EI*_. Δ*γ* - mean value of Pr(*TRCMPLX* = active |∀*ge*_*i*_ evidences) over all runs, *γ* - Pr(*TRCMPLX* = active | all gene evidences), *β* - the assigned cpt value of *TRCMPLX* during initialization of the Bayesian network models and Δ*β* - the deviation in the assigned values of *TRCMPLX* during initialization.

| $\beta$ | $\Delta\beta$ | $\frac{\Delta\beta}{\beta}$ | $\log(1 + \frac{\Delta\beta}{\beta})$ | $\Delta\gamma$ in Normal | $\Delta\gamma$ in Tumor |
| --- | --- | --- | --- | --- | --- |
| 0.8 | 0.1 | 0.125 | 0.117783 | 0.03055427 | 0.09151151 |
| 0.7 | 0.1 | 0.1428571 | 0.1335314 | 0.01423754 | 0.09086427 |
| 0.6 | 0.1 | 0.1666667 | 0.1541507 | 0.004384244 | 0.08052346 |
| 0.5 | 0.1 | 0.2 | 0.1823216 | 0.0005872203 | 0.07294716 |
| 0.8 | 0.1 | 0.125 | 0.117783 | 0.03055427 | 0.09151151 |
| 0.7 | 0.2 | 0.2857143 | 0.2513144 | 0.04479181 | 0.1823758 |
| 0.6 | 0.3 | 0.5 | 0.4054651 | 0.04917605 | 0.2628992 |
| 0.5 | 0.4 | 0.8 | 0.5877867 | 0.04976327 | 0.3358464 |

**Table 2:** Deviation study for model *M*_*PBK*_. Δ*γ* - mean value of Pr(*TRCMPLX* = active |∀*ge*_*i*_ evidences) over all runs, *γ* - Pr(*TRCMPLX* = active | all gene evidences), *β* - the assigned cpt value of *TRCMPLX* during initialization of the Bayesian network models and Δ*β* - the deviation in the assigned values of *TRCMPLX* during initialization.

| $\beta$ | $\Delta\beta$ | $\frac{\Delta\beta}{\beta}$ | $\log(1 + \frac{\Delta\beta}{\beta})$ | $\Delta\gamma$ in Normal | $\Delta\gamma$ in Tumor |
| --- | --- | --- | --- | --- | --- |
| 0.8 | 0.1 | 0.125 | 0.117783 | 0.1400355 | 0.1097089 |
| 0.7 | 0.1 | 0.1428571 | 0.1335314 | 0.06442086 | 0.1877266 |
| 0.6 | 0.1 | 0.1666667 | 0.1541507 | 0.01762791 | 0.06204044 |
| 0.5 | 0.1 | 0.2 | 0.1823216 | 0.01393517 | 0.1718198 |
| 0.8 | 0.1 | 0.125 | 0.117783 | 0.1400355 | 0.1097089 |
| 0.7 | 0.2 | 0.2857143 | 0.2513144 | 0.2044564 | 0.2974356 |
| 0.6 | 0.3 | 0.5 | 0.4054651 | 0.2220843 | 0.359476 |
| 0.5 | 0.4 | 0.8 | 0.5877867 | 0.2360195 | 0.5312958 |

Figures 4, 5, 6 and 7 show the deviations represented in tables 1 and 2. Note that the numbers depicted in the tables are scaled in a nonuniform manner for observational purpose in the figures. Unscaled values are represented under the last two columns on the right of tables 1 and 2. Before reading the graphs, note that red indicates deviation of mean of Pr(*TRCMPLX* = active |∀*ge*_*i*_ evidences) in normal test samples, blue indicates deviation of mean of Pr(*TRCMPLX* = active |∀*ge*_*i*_ evidences) in tumor case, green indicates deviations in Weber’s law and cyan indicates deviations in Bernoulli’s law.

**Figure 4:**
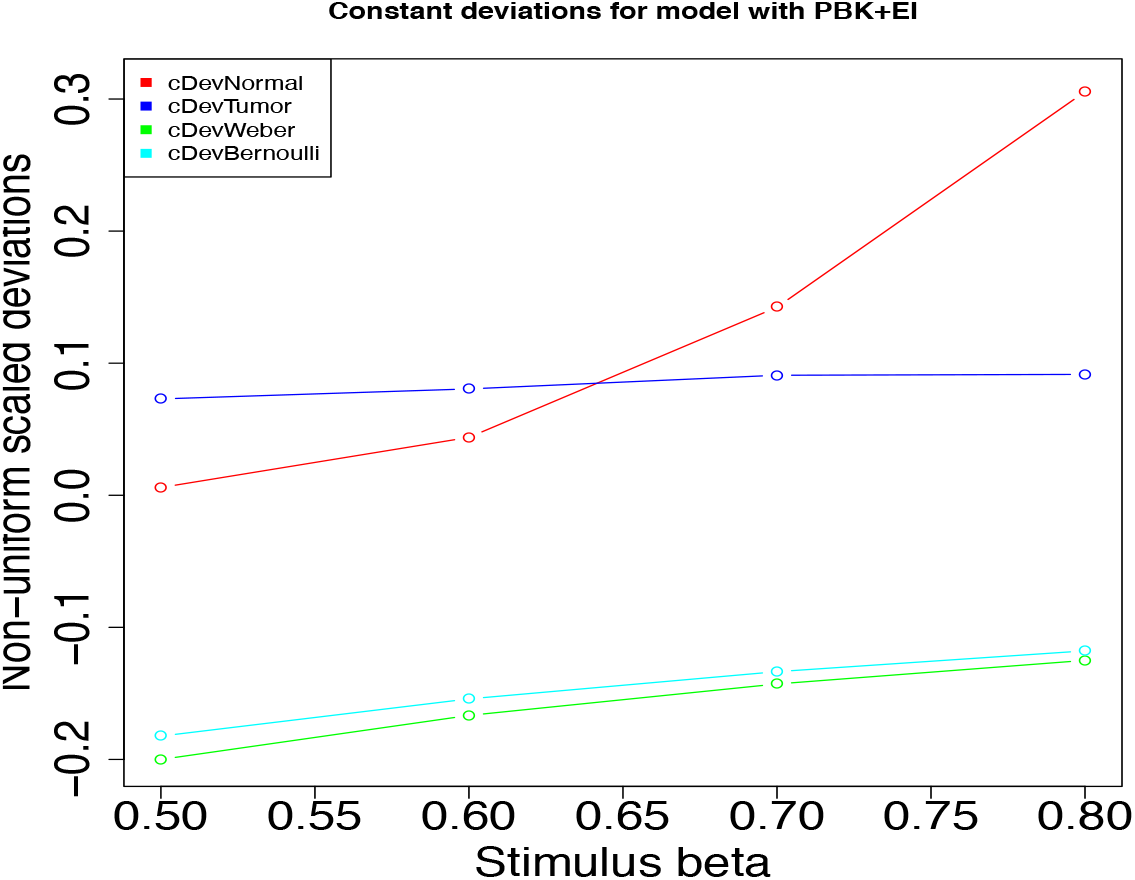
Constant deviations in *β* i.e *ETGN* and corresponding deviations in Pr(*TRCMPLX* = active |∀*ge*_*i*_ evidences) for both normal and tumor test samples. Corresponding Weber and Bernoulli deviations are also recorded. Note that the plots and the y-axis depict scaled deviations to visually analyse the observations. The model used is *M*_*PBK*+*EI*_. Red - constant deviation in Normal, constant deviation in Tumor, Green - constant deviation in Weber’s law, Cyan - constant deviation in Bernoulli’s law.

For the case of contant deviations (figure 4) in model *M*_*PBK*+*EI*_, it was observed that deviations in activation of *TRCMPLX* conditional on gene evidences for the tumor test samples showed a logarithmic behaviour and were directly proportional to the negative of both the Weber’s and Bernoulli’s law. This can be seen by the blue curve almost following the green and cyan curves. For the case of deviations in activation of *TRCMPLX* conditional on gene evidences for the normal test samples showed an exponential behaviour and were proportional to negative of both the Weber’s and Bernoulli’s law. Similar behaviour was observed for all the coloured curves in case of incremental deviations as shown in figure 5. The exponential behaviour for activation of *TRCMPLX* being active conditional on gene evidences correctly supports to the last line of Equation 7 which is the derivation of Weber’s law from Bernoulli’s equation. It actually point to Fechner’s derivation of Weber’s law from logarithmic formulation.

**Figure 5:**
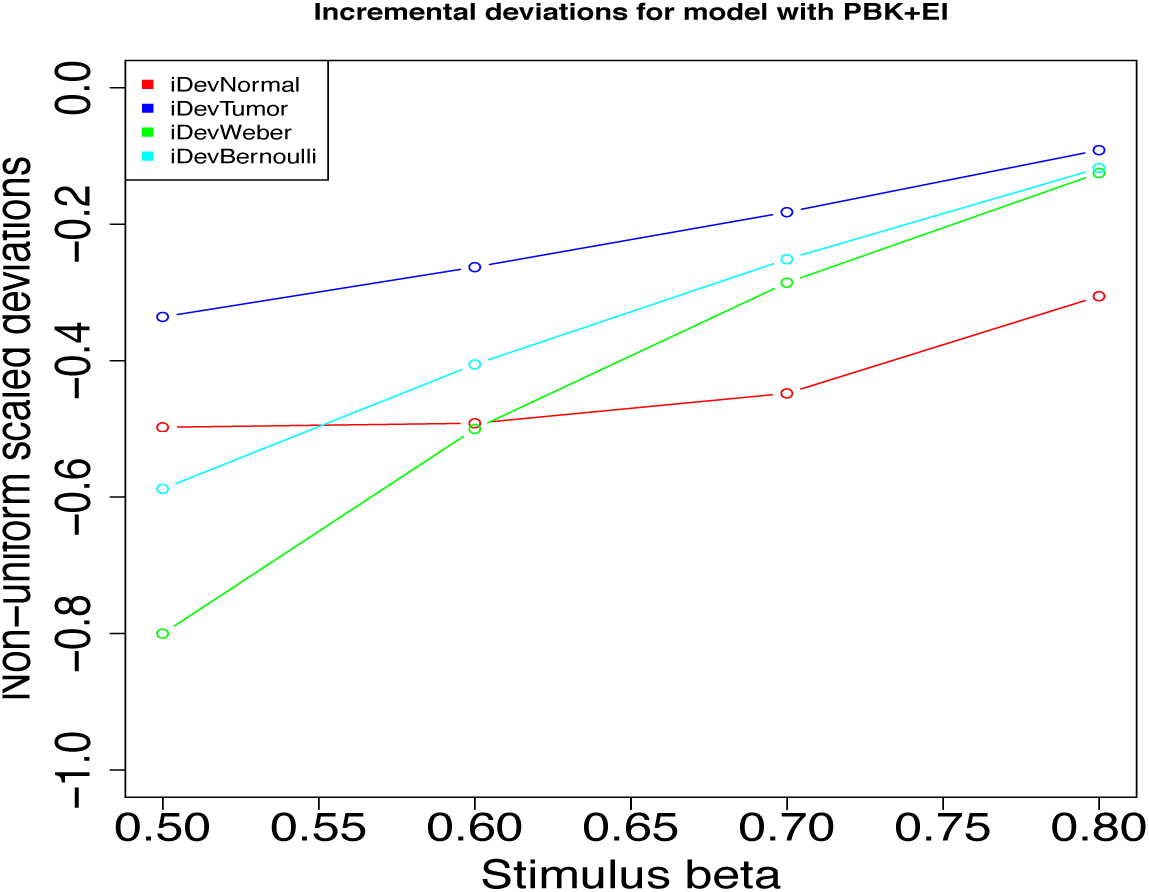
Incremental deviations in *β* i.e *ETGN* and corresponding deviations in Pr(*T RCMP LX* = active |∀*ge*_*i*_ evidences) for both normal and tumor test samples. Corresponding Weber and Bernoulli deviations are also recorded. Note that the plots and the y-axis depict scaled deviations to visually analyse the observations. The model used is *M*_*PBK*+*EI*_. Red - incremental deviation in Normal, incremental deviation in Tumor, Green - incremental deviation in Weber’s law, Cyan - incremental deviation in Bernoulli’s law.

For model *M*_*PBK*_, the above observations do not yield consistent behaviour. In figure 6, for the case of constant deviations, only the deviations in activation of *TRCMPLX* conditional on gene evidences for normal test samples exponential in nature and were found to be directly proportional to the negative of both the Weber’s and Bernoulli’s law. But the deviations in activation of *TRCMPLX* conditional on gene evidences in tumor test samples show noisy behaviour. But this observation is not the case in incremental deviations for the same model. For the case of incremental deviations as represented in figure 7, the deviations in activation of *TRCMPLX* conditional on gene evidences is directly proportional to both the Weber’s and Bernoulli’s law. The figure actually represent the plots with inverted values i.e negative values. A primary reason for this behaviour might be that *M*_*PBK*_ does not capture and constrain the network as much as *M*_*PBK*+*EI*_ which include epigenetic information. This inclusion of heterogeneous information adds more value to the biologically inspired network and reveals the hidden natural laws occurring in the signaling pathway in both normal and tumor cases.

**Figure 6:**
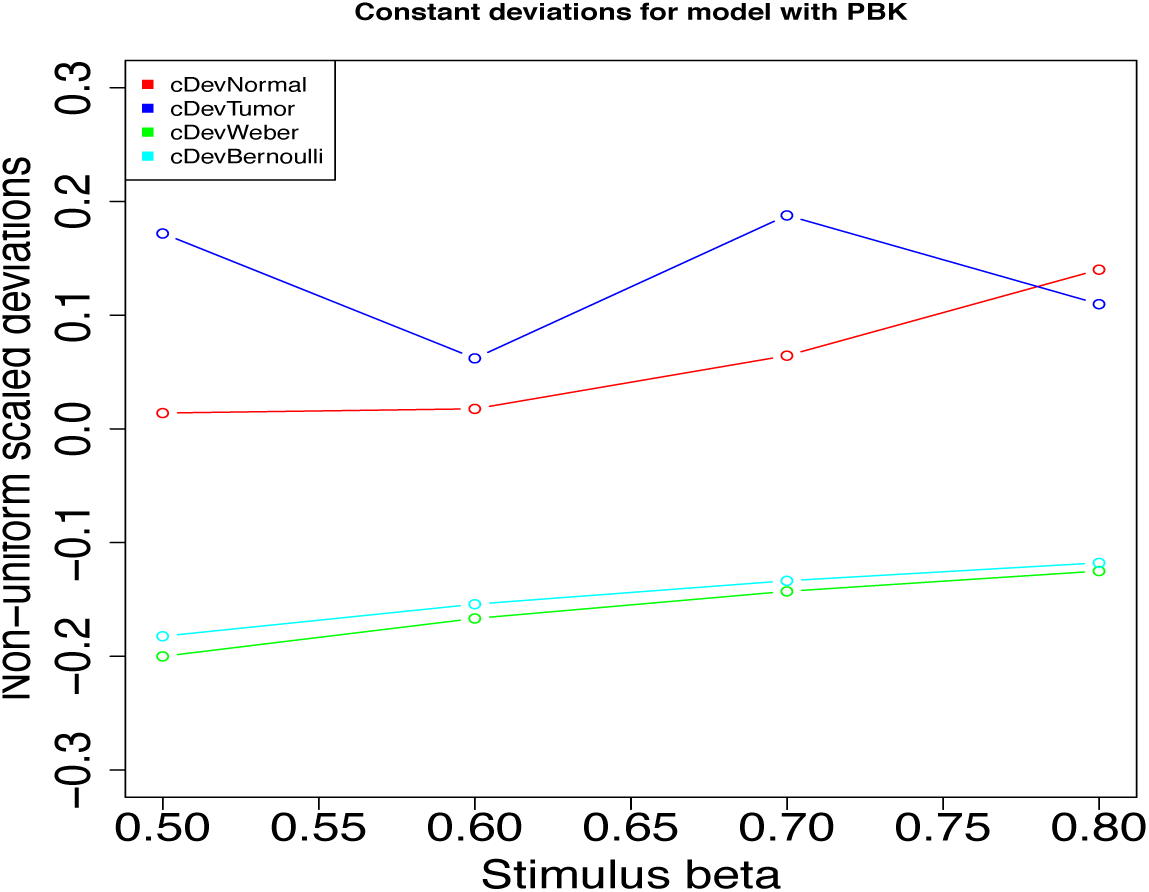
Constant deviations in *β* i.e *ETGN* and corresponding deviations in Pr(*TRCMPLX* = active |∀ *ge*_*i*_ evidences) for both normal and tumor test samples. Corresponding Weber and Bernoulli deviations are also recorded. Note that the plots and the y-axis depict scaled deviations to visually analyse the observations. The model used is *M*_*PBK*_. Red - constant deviation in Normal, constant deviation in Tumor, Green - constant deviation in Weber’s law, Cyan - constant deviation in Bernoulli’s law.

**Figure 7:**
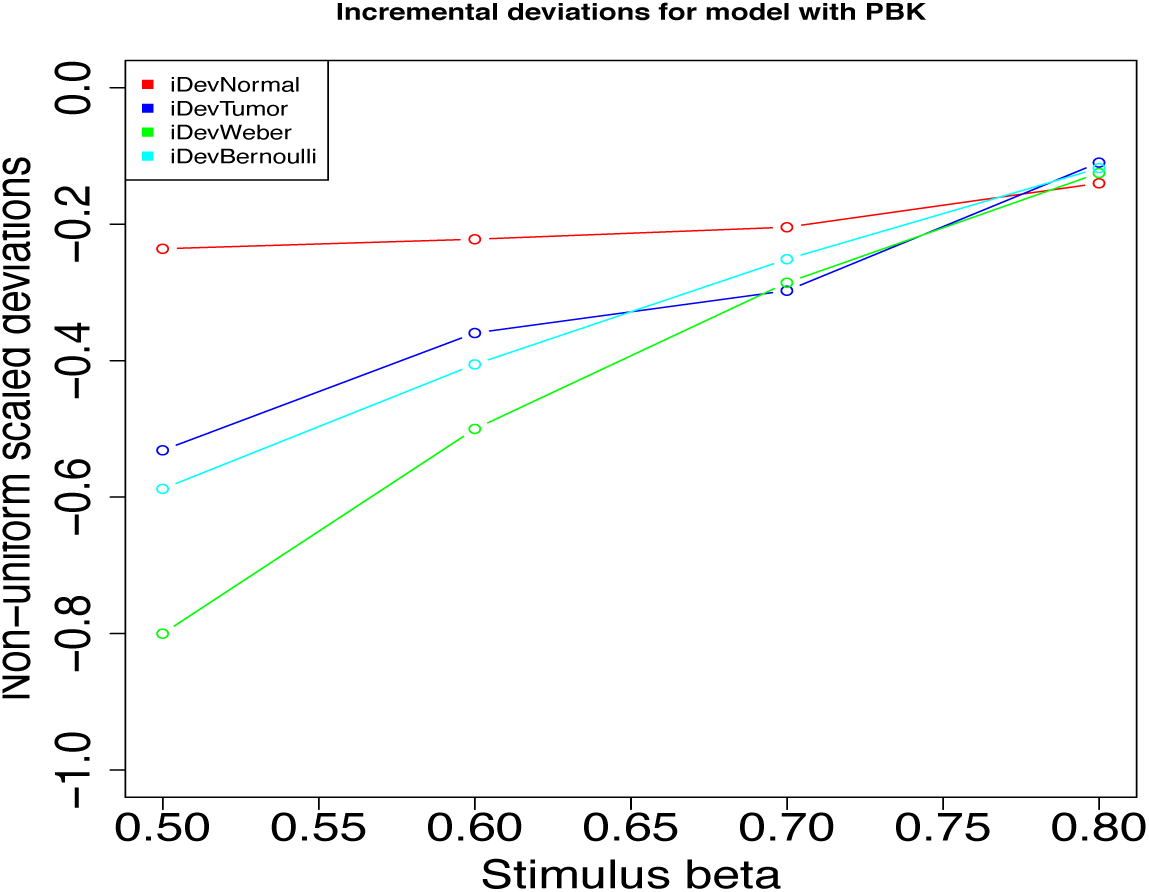
Incremental deviations in *β* i.e *ETGN* and corresponding deviations in Pr(*T RCMP LX* = active |∀ *ge*_*i*_ evidences) for both normal and tumor test samples. Corresponding Weber and Bernoulli deviations are also recorded. Note that the plots and the y-axis depict scaled deviations to visually analyse the observations. The model used is *M*_*PBK*_. Red - incremental deviation in Normal, incremental deviation in Tumor, Green - incremental deviation in Weber’s law, Cyan - incremental deviation in Bernoulli’s law.

### 4.2 Intuition behind the curve behaviour

Lastly, the intuitive idea behind the behaviour of the curves generated from constant deviation in table 1 is as follows. It is expected that Pr(*TRCMPLX* = active*|*all gene evidences) is low (high) in the case of Normal (Tumor) samples. The change ΔPr(*TRCMPLX* = active*|*all gene evidences) jumps by power of 10 as the *β* values change from 50% to 90% in Normal cases. It can be observed from the table that there are low deviations in Pr(*TRCMPLX* = active | all gene evidences) when *β* is low i.e the effect of transcription complex is low and high deviations in Pr(*TRCMPLX* = active | all gene evidences) when *β* is high i.e the effect of transcription complex is high. But it should be noted that the deviations still tend to be small. This implies that the *TRCMPLX* is switched off at a constant rate. Thus changes in *β* leads to exponential curves as in the formulation 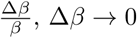, Δ*β →* 0 and *β → ∞*.

In tumor cases, ΔPr(*TRCMPLX* = active | all gene evidences) behaves near to logarithmic curve as *β* increases from 50% to 90%. The deviations increase in a slow monotonic way as *β* increases. Finally, the ratio 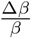 shows monotonically increasing behaviour as Δ*β* increases proportionally with *β*. This means that in tumor samples the rate of transcription increases or the effect of rate of transcription complex increases monotonically as *β* increases. This points to the fold change in *β*-*catenin* concentration that might be influencing the transcription rate of the transcription complex. In normal case, the *β*-*catenin* concentration remains constant. Due to this, changes in the rate of transcription by the transcription complex might remains constant and near to zero. Change in *β* values that is the change in initialization of cpt values of transcription complex causes the logarithmic curve in deviations of prediction of transcription complex.

Finally, these observations present a crude yet important picture regarding the downstream transcriptional behaviour of signaling pathway in case of colorectal cancer. Though the current model does not measure the fold changes in the concentration levels of *β*-*catenin*, it can help in measuring the deviations in activity of the transcription complex conditional on the gene evidences by observing the deviations in the strength of belief assigned as priors in the probability tables of the node representing the transcription complex of the network. Thus sensitivity analysis facilitates in observing such natural phenomena at computational level. In context of the work by Goentoro and Kirschner (2009), the presented results are crude in terms of static observations yet they show corresponding behaviour of transcriptional activity in terms of psychophysical laws. Further investigations using dynamic models might reveal more information in comparison to the static models used in Sinha (2014). The observations presented here bolster the existence of behavioural phenomena in terms of logarithmic laws.

## 5. Preservation of gene gene interactions

The second part of this study was to find interactions between two genes by observing the conditional probability of activation status of one gene given the evidence of another gene. Let *g* be a gene. To obtain the results, two steps need to be executed in a serial manner. The first step is to construct gene gene interactions based on the available conditional probabilities denoted by Pr(*g*_*i*_ = active/repressed | *g*_*k*_ evidence) ∀ *k* genes. The conditional probabilities are inferred using the junction tree algorithm that employs two-pass message passing scheme. Example code and implementations of the same can be found in Murphy et al. (2001). The steps for constructing the gene gene interactions based on these conditional probabilities are documented in the Appendix. The second step is to infer gene gene interaction network based purely on reversible interactions. Note that networks are inferred for gene evidences using normal and tumor test samples separately.

Finally, once the interaction network is ready, the computational empirical estimates for deviations in gene-gene interaction is recorded and observation on the prevalence of psychophysical laws in these interactions is discussed. An important point that needs to be kept in mind is that the inferred interaction network differs based on the choice of the threshold involved (which is a computational issue) but the underlying psychophysical laws remain unchanged (which is a natural phenomena irrespective of the components involved). Thus the while reading the observations on the psychophysical laws, readers must not get confused regarding plots made for interactions from different networks.

### 5.1. Constructing gene-gene interactions

Gene interactions are constructed by labeling the inferred conditional probability of activation of *g*_*j*_ given the state of *g*_*i*_, for all *j′ s&i′ s*. Here labels refer to assigning *<>* for an activated gene and *|* for repressed gene. Thus the following possible combinations can be inferred - *g*_*j*_ *<>* − *<> g*_*i*_, *g*_*j*_*|* − *|g*_*i*_, *g*_*j*_ *<>* −*|g*_*i*_ and *g*_*j*_*|*− *<> g*_*i*_. Note that all interactions are basically depicting the degree of belief in the state of *g*_*i*_ given or conditional on *g*_*j*_ i.e Pr(*g*_*i*_*|g*_*j*_). The label related to *g*_*i*_ is derived by discretizing Pr(*g*_*i*_*|g*_*j*_) with respect to a weighted mean or the arbitrary value of 0.5. In any interaction, the label associated with *g*_*j*_ is the evidence and the label associated with *g*_*i*_ is the predicted conditional probability. Thus there will always exist two way interactions corresponding to Pr(*g*_*j*_ *| g*_*i*_) and Pr(*g*_*i*_ *| g*_*j*_) in a Normal case. Similar interactions can be inferred for the Tumor case. Which interactions to select is based on criteria of reversibility and duplication, which is addressed later. To reiterate a final note regarding the interactions - the inferred interactions differ based on the choice of the threshold involved (which is a computational issue) but if prevalent, the underlying psychophysical laws remain unchanged (which is a natural phenomena irrespective of the components involved).

The network obtained by using an arbitrary value like 0.5 for labeling the gene interactions is different from those obtained using a weighted mean. There are advantages of choosing the weighted mean of the training labels for each gene - • Each gene has an individual threshold that is different from the other as the expression values are different and the discretization used to estimate a particular threshold is based on the median value of the training data for that particular gene under consideration. • The weighted mean assigns appropriate weights to the labels under consideration rather than assigning equal weights which might not represent the actual threshold. • Due to the properties mentioned in the second point, it might be expected that the weighted mean generates a sparse network in comparison to that generated using an arbitrary value of 0.5. • Finally, the weighted mean could reveal interactions between two genes that might be happening at different stages of time. Even though using a static model, capturing such intricate interactions is possible as will be seen later.

There is a formulation for weighted means, but the computation of the weighted mean for training samples belonging to Normal and Tumor is done separately. The separate formulations are given below -

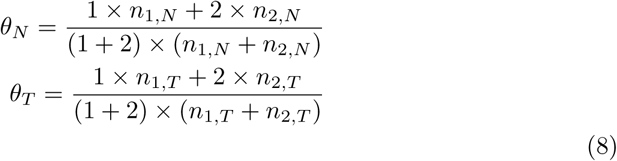

were, *n*_1*,N*_ is the number of Normal training samples with label 1, *n*_2*,N*_ is the number of Normal training samples with label 2, *n*_1*,T*_ is the number of Tumor training samples with label 1 and *n*_1*,N*_ is the number of Tumor training samples with label 2. Note that the sample labels (i,e evidence of gene expression) were discretized to passive or 1 (active or 2).

Based on the steps described in Appendix, for each gene a matrix is obtained that shows the statistics of how the status of a gene is affected conditional on the individual evidences of the remaining genes. Also, for each of the *i*^*th*^ gene the averaged 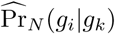 is also stored in vector *PggN*. Same is done for tumor cases. These two vectors are later used to test the veracity of existence of psychophysical laws in gene-gene interaction network. Table 3 represents one such tabulation for gene *SFRP* 3. For all runs and all test samples, the following was tabulated in table 3: *aa*_*N*_ - *SFRP* 3 is active (a) when a gene is active (a) in Normal (N) sample, *ar*_*N*_ - *SFRP* 3 is active (a) when a gene is repressed (r) in Normal (N) sample, *ra*_*N*_ - *SFRP* 3 is repressed (r) when a gene is active (a) in Normal (N) sample, *rr*_*N*_ - *SFRP* 3 is repressed (r) when a gene is repressed (r) in Normal (N) sample, *aa*_*T*_ - *SFRP* 3 is active (a) when a gene is active (a) in Tumor (T) sample, *ar*_*T*_ - *SFRP* 3 is active (a) when a gene is repressed (r) in Tumor (T) sample, *pa*_*T*_ - *SFRP* 3 is repressed (r) when a gene is active (a) in Tumor (T) sample, *gg*_*IN*_ - interaction of *SFRP* 3 given the gene evidence based on majority voting among *aa*_*N*_, *ar*_*N*_, *ra*_*N*_ and *rr*_*N*_ and finally, *gg*_*IT*_ - interaction of *SFRP* 3 given the gene evidence based on majority voting among *aa*_*T*_, *ar*_*T*_, *ra*_*T*_ and *rr*_*T*_. The highest score among *aa*_*N*_, *ar*_*N*_, *ra*_*N*_ and *rr*_*N*_ (*aa*_*T*_, *ar*_*T*_, *ra*_*T*_ and *rr*_*T*_) confirms the relation between genes using Normal (Tumor) samples. Activation (repression) for *SFRP* 3 is based on discretizing the predicted conditional probability Pr(*SFRP* 3 = active*|g*_*j*_ evidence) as *≥ θ* (*< θ*). Activation (repression) for a particular gene evidence *g*_*j*_ is done using discrete evidence. In table 3, under the columns *gg*_*IN*_ and *gg*_*IT*_, *<>* implies the gene is active and | implies the gene is repressed or passive.

**Table 3:**
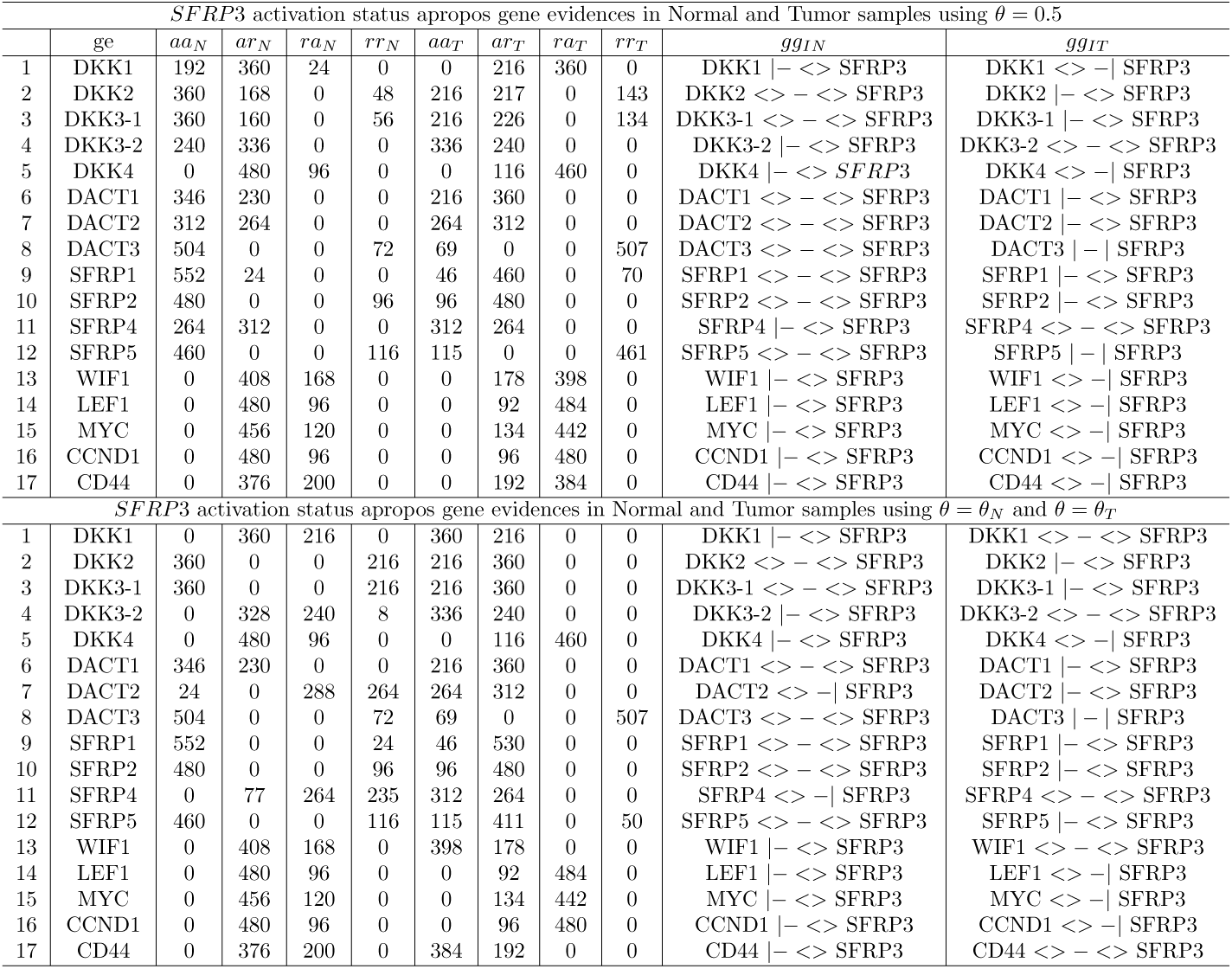
*SFRP* 3 activation status in test samples conditional on status of individual gene activation (represented by evidence in test data) in Normal and Tumor samples. Measurements are taken over summation of all predicted values across the different runs of the 2-Hold out experiment. Here the notations denote the following: a - active, p - passive, N - Normal, T - Tumor, *gg*_*IN*_ - gene-gene interaction with Normal, *gg*_*IT*_ - gene-gene interaction with Tumor, *<>* - active and *|* - repressed.

**Gene-gene interaction network when** *θ* = 0.5

Considering only reversible interactions, in table 3 it was found that evidence for *DKK*1 and *DKK*4 show similar repression behaviour as the standard genes *WIF* 1, *LEF* 1, *MY C*, *CCND*1 and *CD*44 in Normal (Tumor) test samples. Only, *SFRP* 5 and *DACT* 3 in Normal (Tumor) test samples shows activation (repression). Conditional on the observed activation status of the genes mentioned above, *SFRP* 3 shows activated (repressed) state in Normal (Tumor) test samples. *SFRP* 3 showed behaviour similar to *SFRP* 1*/*2*/*5. Since it is known that the activation status of the latter is influenced by epigenetic factors, *SFRP* 3 might also be influenced by epigenetic factors.

Irreversible interactions present in table 3 are deleted as they do not provide concrete information regarding the functional roles of the genes in normal and tumor cases. This attributes to one of the following facts (1) noise that corrupts prediction values as can be seen in the columns of *aa*_*N*_ (*aa*_*T*_), *ar*_*N*_ (*ar*_*T*_), *ra*_*N*_ (*ra*_*T*_) and *rr*_*N*_ (*rr*_*T*_) or (2) other multiple genes might be interacting along with *SFRP* 3 in a combined manner and it is not possible to decipher the relation between *SFRP* 3 and other genes. This calls for investigation of prediction of *SFRP* 3 status conditional on joint evidences of two or more genes (a combinatorial problem with a search space order of 2^17^ − 17, which excludes 17 cases of individual gene evidences which have already been considered here). Incorporating multiple gene evidences is not a problem while using Bayesian network models as they are designed to compute conditional probabilities given joint evidences also (except at the cost of high computational time).

It is evident that an arbitrary value of *θ* = 0.5 will not generate appropriate networks. This is due to the fact that 0.5 does not encode the biological knowledge of thresholding while using discretization. To over come this, a weighted mean is employed as shown below.

**Gene-gene interaction network when 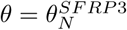**

While employing the weighted mean as the threshold to discretize Pr(*SFRP* 3 = active | *g*_*j*_ evidence), the *SFRP* 3 gene evidences that constitutes the test data are used. See step 5.a.iii in Appendix. Note that the test evidences for *SFRP* 3 are used for two purpose (1) to discretize Pr(*SFRP* 3 = active | *g*_*j*_ evidence) as discussed above and (2) to compute the probability of activation status of another gene conditional on evidence for *SRFP* 3, i.e Pr(*g*_*j*_ = active | *SFRP* 3 evidence). Why to use test evidence or labels to compute weighted mean? Since the test evidence for a gene (i.e the discretized label) has been derived using the median computed on the corresponding training data for the same gene, it absolutely fine to use the discretized test labels to further compute the weighted mean. This is because the median of gene expression is a value which is much higher than the probability value of 1 and cannot be used to discretize a predicted conditional probability value. Also, estimating the density estimates from a small population of gene expression values has its own weakness. To converge on a plausible realistic value the discretized test samples can be used to estimate a weighted mean which represents the summary of the distribution of the discretized values. This weighted mean of *SFRP* 3 test samples then discretizes Pr(*SFRP* 3 = active | *g*_*j*_ evidence) according to the inherently represented summary. More realistic estimates like kernel density estimates could also be used.

In comparison to the interactions derived using *θ* = 0.5 in table 3, it was found that a more restricted list of *DKK*4, *DACT* − 2*/*3, *LEF* 1, *MY C* and *CCND*1 showed reversible behaviour with *SFRP* 3 using the weighted mean. This reduction in the reversible interactions is due to the fact that the weighted mean carries an idiosyncrasy of the test label data distribution and is more restricted in comparison to the use of 0.5 value that was arbitrarily chosen. Finally, using the proposed weighted mean reveals more than one interaction between two genes. These interactions point to important hidden biological phenomena that require further investigation in the form of wet lab experiments and the ensuing in silico analysis. It also points to the fact that a particular gene may be showing different behaviour at different times in the network while interacting with multiple genes. An example of this will be addressed later. Again, dynamic models will bring more clarity to the picture. Table 3 shows these interactions using *θ ∈ {*0.5*, θ*_*N*_*, θ*_*T*_ *}*.

### 5.2. Inferring gene-gene interaction network

Next, after the construction of gene-gene interactions, it is necessary to infer the network. The inference of the estimated gene-gene interactions network is based on explicitly reversible roles in Normal and Tumor test samples. This means that only those interactions are selected which show the following property - *g*_*j*_ *<>* − *<> g*_*i*_ in Normal if and only if *g*_*j*_*|* − *|g*_*i*_ in Tumor, *g*_*j*_ *<>* −*|g*_*i*_ in Normal if and only if *g*_*j*_*|*− *<> g*_*i*_ in Tumor, *g*_*j*_*|*− *<> g*_*i*_ in Normal if and only if *g*_*j*_ *<>* −*|g*_*i*_ in Tumor and finally, *g*_*j*_*|* − *|g*_*i*_ in Normal if and only if *g*_*j*_ *<>* − *<> g*_*i*_. This restricts the network to only reversible gene-gene interactions in Normal and Tumor cases. Note that an interaction *g*_*j*_*ℐ ℛg*_*i*_ (*g*_*i*_*ℐ ℛg*_*j*_) is depicted by Pr(*g*_*i*_ | *g*_*j*_) (Pr(*g*_*j*_ | *g*_*i*_)).

Reversibility helps in tracking the behaviour of gene-gene interaction in both normal and tumor case simultaneously and thus give more weight to confirmatory results. Irreversible reactions here mean that the state of activation of a gene in both normal and tumor sample remains invariant given the evidence of the other gene in the gene-gene interaction. This helps in eliminating the interactions that might not be happening at all from biological perspective. To confirm the computational results wet lab experiments are needed. See table 3 for reversible and irreversible interactions.

Next, duplicate interactions are removed from the network for normal samples. This is repeated for the network based on tumor samples also. This is achieved by removing one of the interactions from the following pairs (*g*_*j*_ *<>* − *<> g*_*i*_ and *g*_*i*_ *<>* − *<> g*_*j*_), (*g*_*j*_ *<>* −*|g*_*i*_ and *g*_*i*_*|*− *<> g*_*j*_), (*g*_*j*_*|*− *<> g*_*i*_ and *g*_*i*_ *<>* −|*g*_*j*_) and (*g*_*j*_ |−| *g*_*i*_ and *g*_*i*_ | − |*g*_*j*_). This process is done to remove redundant interactions that are recorded via steps mentioned in construction of gene-gene interaction network. Figure 8 shows one such network after complete network construction, interaction labeling, consideration of reversible interactions and removal of duplicate interactions using Normal test samples with ETGN of 90% in *M*_*PBK*+*EI*_. For the case of Tumor test samples with ETGN 90% in *M*_*PBK*+*EI*_, only the reversal of interactions need to be done. Table 4 and 5 represents these interactions in figures 8 and 9 in a tabulated form, respectively.

**Table 4:**
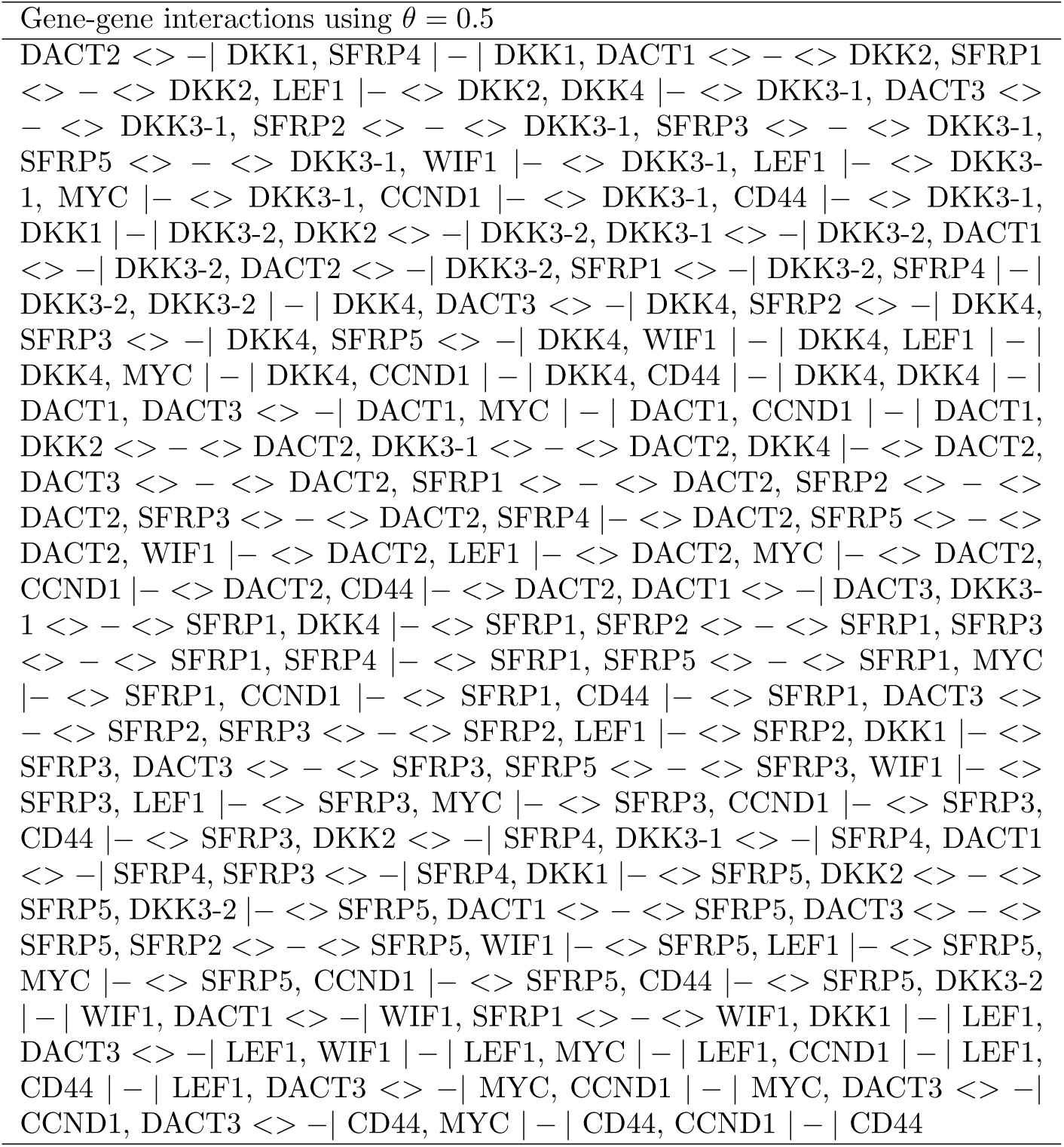
Tabulated gene gene interactions of figure 8 using *M*_*PBK*+*EI*_ obtained in case of Normal samples. Here, the symbols represent the following - *<>* activation and | repression/suppression. Note that for Tumor cases, the interaction roles were found to be reversed, ie. *<>* −*|* in normal became *|*− *<>* in tumor, *|*− *<>* in normal became *<>* −*|* in tumor, *<>* − *<>* in normal became *|* − *|* in tumor and *|* − *|* in normal became *<>* − *<>* in tumor.

**Table 5:**
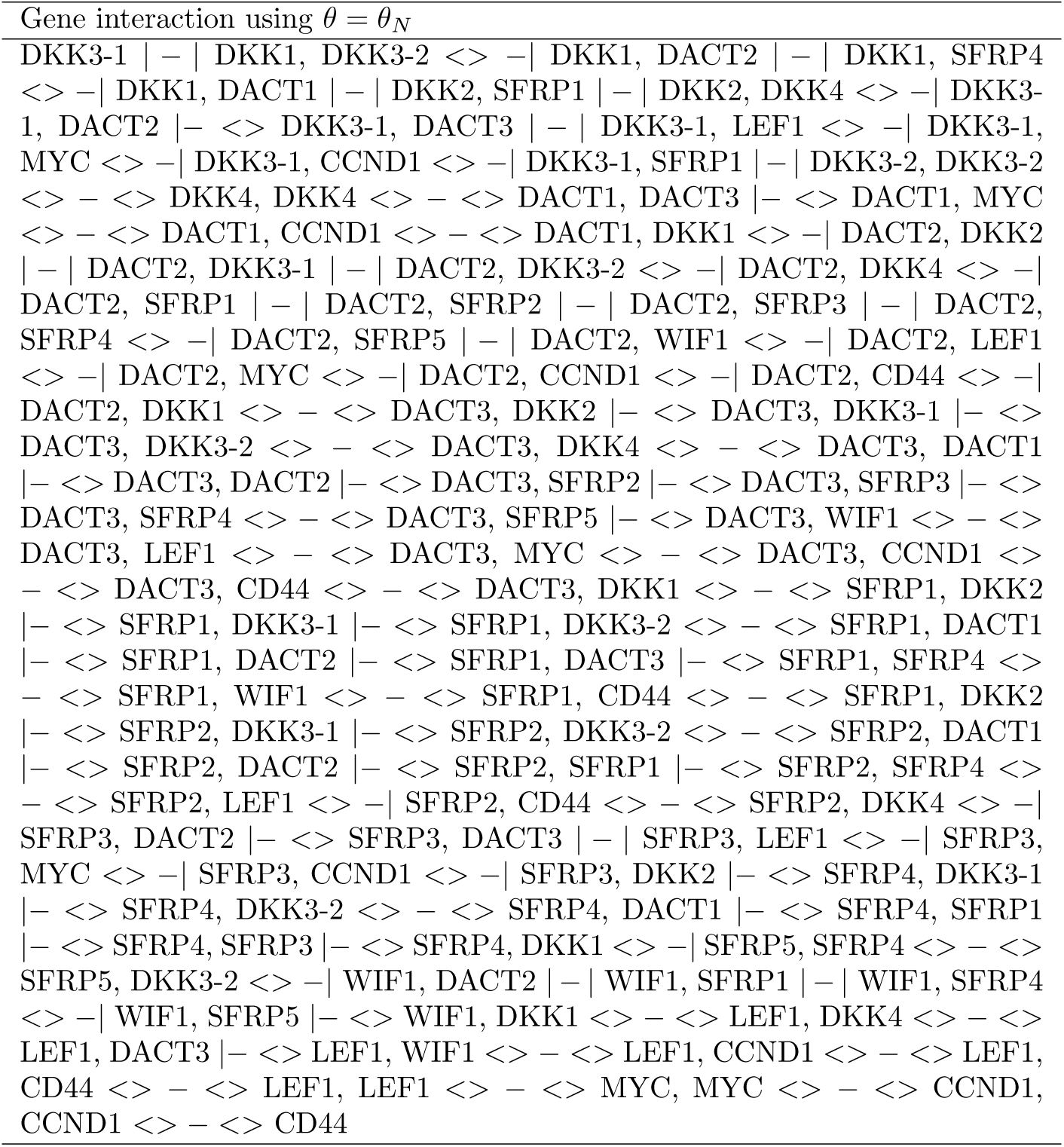
Tabulated gene gene interactions of figure 9 using *M*_*PBK*+*EI*_ obtained in case of Normal samples. Here, the symbols represent the following - *<>* activation and | repression/suppression. Note that for Tumor cases, the interaction roles were found to be reversed, ie. *<>* −*|* in normal became *|*− *<>* in tumor, *|*− *<>* in normal became *<>* −*|* in tumor, *<>* − *<>* in normal became *|* − *|* in tumor and *|* − *|* in normal became *<>* − *<>* in tumor.

**Figure 8:**
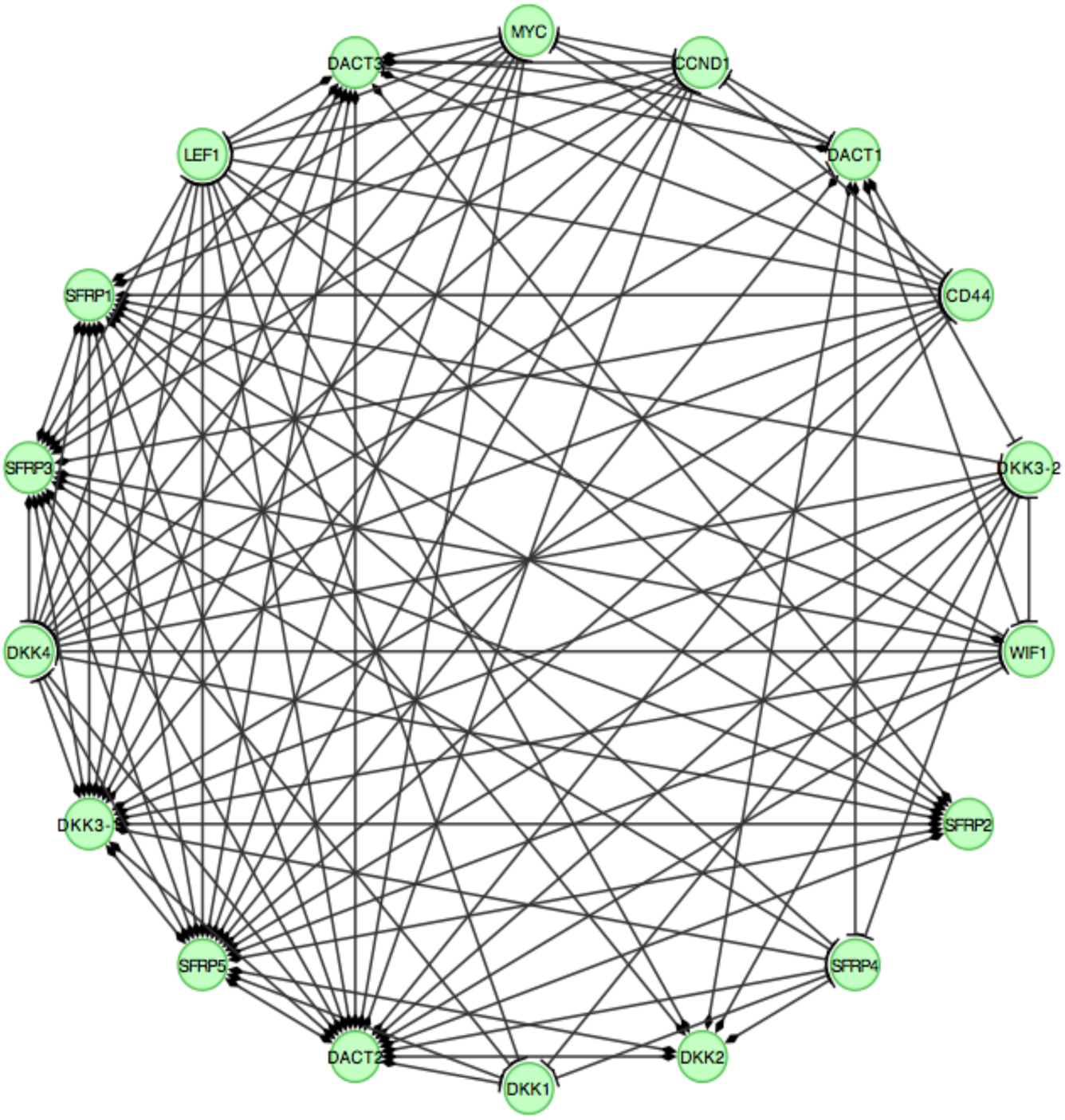
Gene gene interactions for normal case while using *M*_*PBK*+*EI*_ with *θ* = 0.5. Note that the effect of initialized cpt for *TRCMPLX* is 90% in tumorous case and 10% in normal case. Diamond *<>* means activation and straight bar *|* means repression.

**Figure 9:**
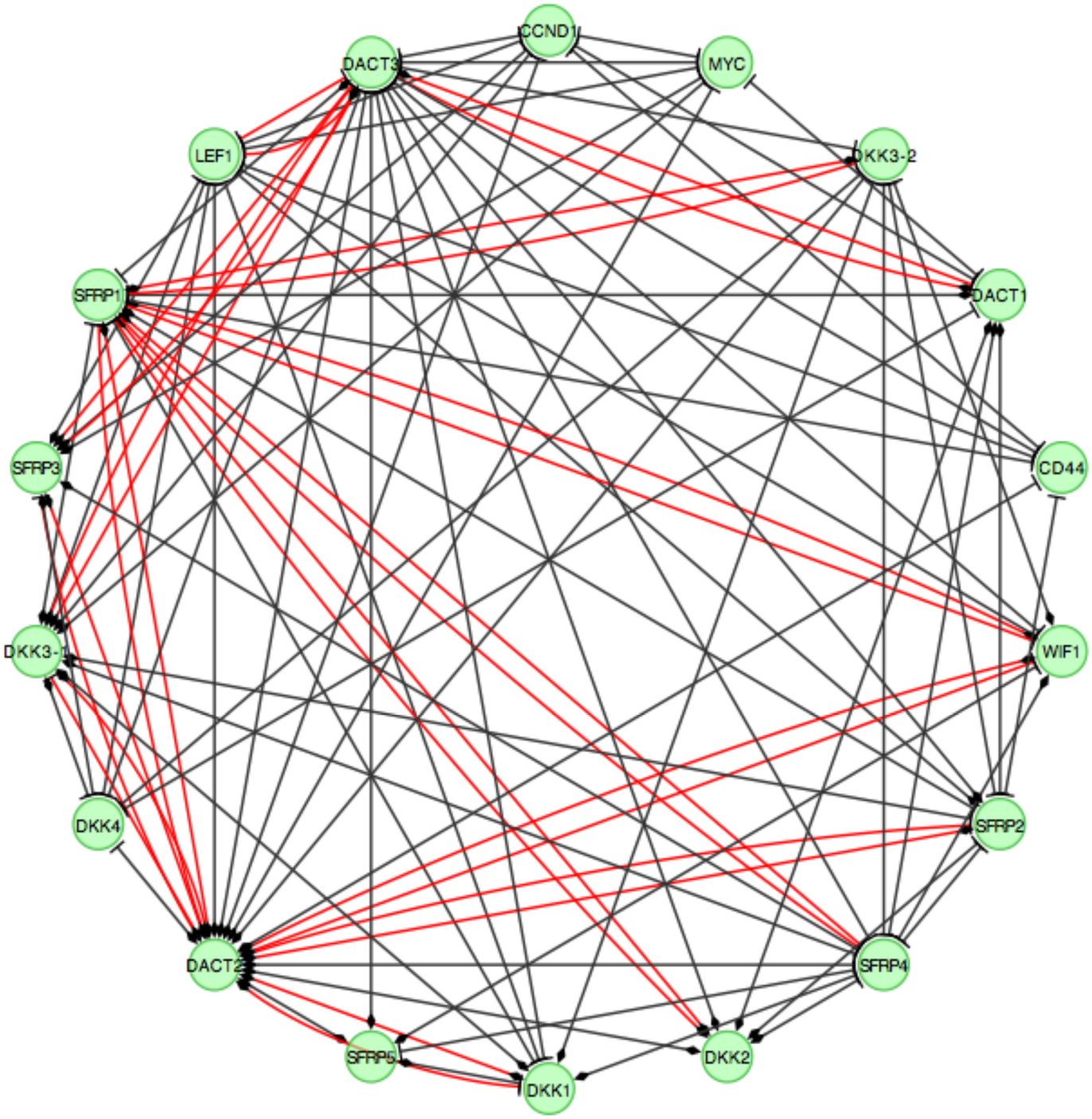
Gene gene interactions for normal case while using *M* _*PBK*+*EI*_ with *θ* = *θ*_*N*_. Note that the effect of initialized cpt for *TRCMPLX* is 90% in tumorous case and 10% in normal case. Diamond *<>* means activation and straight bar *|* means repression.

Finally, different networks were generated by varying the effect of *TRCMPLX* (ETGN) and compared for the normal test samples. Table 6 represents the different interactions that were preserved in network from ETGN 90% with respect to networks obtained from ETGN with values of 80%, 70%, 60% and 50%. It was found that most of the genetic interactions depicted in figures 8 and 9 were found to be preserved across the different variations in ETGN as shown in table 6. Out of the total *n* genes which construct a fully connected graph of 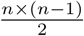, it was observed that lesser number of interconnections were preserved. This preservation indicates towards the robustness of the genetic contributions in the Wnt signaling pathway in both normal and tumor test samples. Note that these observations are made from static models and dynamic models might reveal greater information.

**Table 6:**
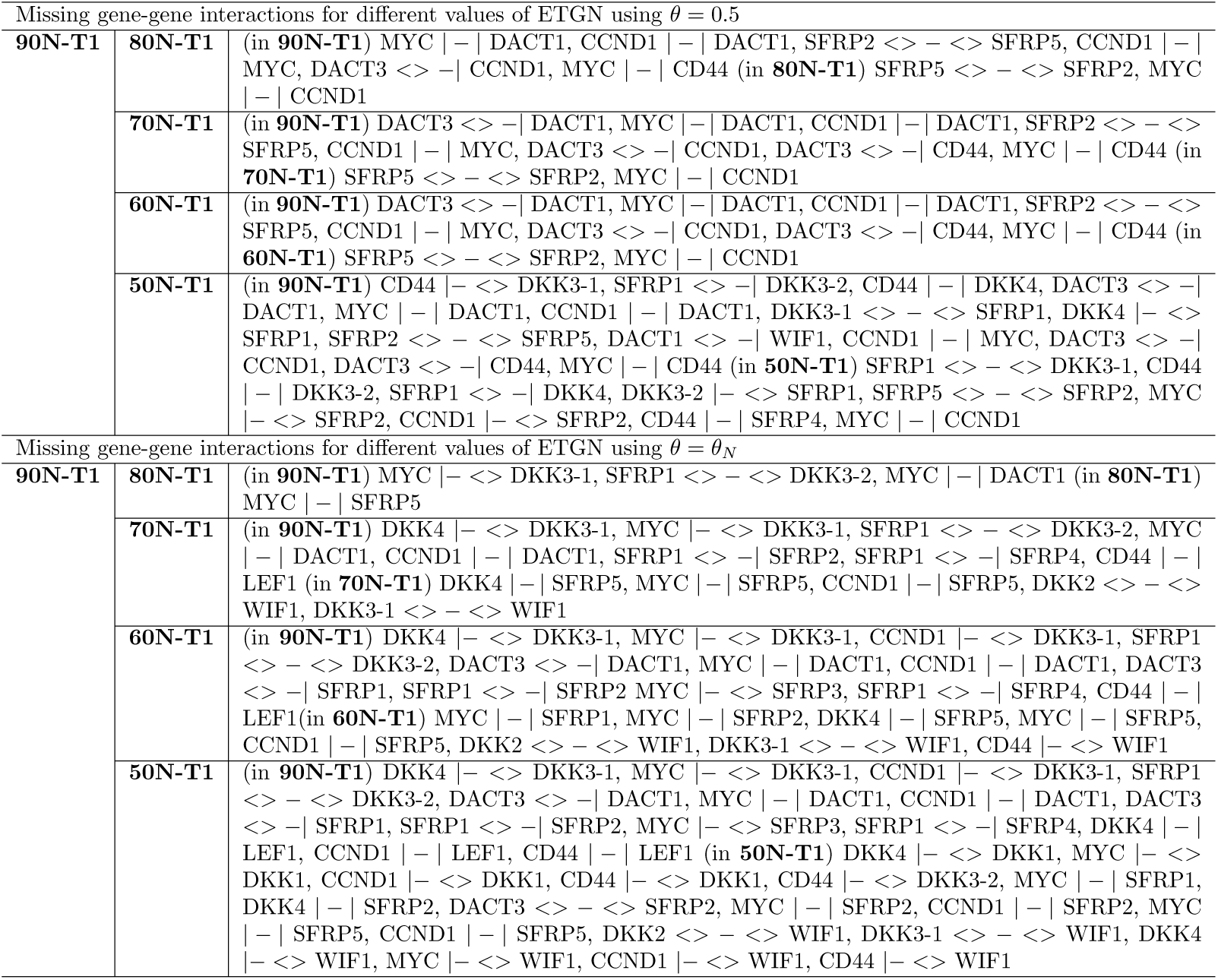
Tabulated missing gene gene interactions of figure 8 and 9 using *M*_*PBK*+*EI*_ obtained in case of Normal samples. Interactions found in Normal samples with 80%, 70%, 60% and 50% effect that are not found with 90% and vice versa have been recorded. Here, the symbols represent the following - *<>* activation and *|* repression/suppression. Note that for Tumor cases, the interaction roles were found to be reversed, ie. *<>* −*|* in normal became *|*− *<>* in tumor, *|*− *<>* in normal became *<>* −*|* in tumor, *<>* − *<>* in normal became *|* − *|* in tumor and *|* − *|* in normal became *<>* − *<>* in tumor.

## 6. Results and discussion on observations 2 & 3

### 6.1. Logarithmic-power deviations in prediction of gene-gene interactions

In the previous section, it was found that some of the interactions remain preserved as there was change in the affect of transcription complex. The first observation of this work was that deviations in the activity of the transcription complex followed a logarithmic-power psychophysical law. The manifestation of these laws at transcriptional levels can be attributed to the fold changes in *β*-*catenin* levels and the prevalence of Weber’s law observed by Goentoro and Kirschner (2009). In this perspective, it would be interesting to observe if these laws are prevalent among the gene-gene interactions in the network or not.

**Case:** *<>* −| **or** |−*<>* **with** *θ* = *θ*_*N*_

In Sinha (2014), the unknown behaviour of *SFRP* 3 in the Wnt pathway has been revealed slightly using computational causal inference. In figure 8, *SFRP* 3 shows preservation in the network and it’s interaction with other genetic factors involved in the model proposed in Sinha (2014) has been depicted. In one such paired interaction between *SFRP* 3 and *MY C*, *SFRP* 3 showed activation (repression) and *MY C* showed repression (activation) in normal (tumor) samples. As the change in the effect of transcription complex was induced by changing the initially assigned cpt values for *TRCMPLX* node, the deviations in the prediction of the gene-gene interaction network was observed to follow the logarithmic-power law crudely. What this means is that deviations or fold changes might also be prevalent at the gene-gene interaction level due to the upstream fold changes in *β*-*catenin* that induces transcriptional activity. More specifically, the deviation in the joint interaction that is represented by the degree of belief via the conditional probability of status of one gene given the evidence or activation status regarding another gene i.e Pr(*g*_*i*_ | *g*_*j*_), is influenced by the fold changes upstream of the pathway and thus exhibit similar psychophysical laws.

Table 7 and 8 show these deviations in the prediction of the interactions for both the normal and the tumor cases. The tables show how deviations are affected when the changes in the effect of the transcription complex are done at constant and incremental rate. To summarize the results in these tables, graphs were plotted in figures 10 for Pr(*SFRP* 3 | *MY C*) (constant deviations), 11 for Pr(*MY C* | *SFRP* 3) (constant deviations), 12 for Pr(*SFRP* 3 | *MY C*) (incremental deviations) and 11 for Pr(*MY C* | *SFRP* 3) (incremental deviations).

**Table 7:** Deviation study for Pr(*SFRP* 3*|MY C*) and Pr(*MY C|SFRP* 3) for normal case

| $\beta$ | $\Delta\beta$ | $\frac{\Delta\beta}{\beta}$ | $\log(1 + \frac{\Delta\beta}{\beta})$ | $\Delta\gamma$<br>$\Pr(SFRP3\vert MYC)$ | $\Delta\gamma$<br>$\Pr(MYC\vert SFRP3)$ |
| --- | --- | --- | --- | --- | --- |
| 0.8 | 0.1 | 0.125 | 0.117783 | 0.003014287 | 0.00324456 |
| 0.7 | 0.1 | 0.1428571 | 0.1335314 | 0.002766111 | 0.00324456 |
| 0.6 | 0.1 | 0.1666667 | 0.1541507 | 0.002504868 | 0.00324456 |
| 0.5 | 0.1 | 0.2 | 0.1823216 | 0.002228110 | 0.00324456 |
| 0.8 | 0.1 | 0.125 | 0.117783 | 0.010513376 | 0.01297824 |
| 0.7 | 0.2 | 0.2857143 | 0.2513144 | 0.007499089 | 0.00973368 |
| 0.6 | 0.3 | 0.5 | 0.4054651 | 0.004732978 | 0.00648912 |
| 0.5 | 0.4 | 0.8 | 0.5877867 | 0.002228110 | 0.00324456 |

**Table 8:** Deviation study for Pr(*SFRP* 3*|MY C*) and Pr(*MY C|SFRP* 3) for tumor case

| $\beta$ | $\Delta\beta$ | $\frac{\Delta\beta}{\beta}$ | $\log(1 + \frac{\Delta\beta}{\beta})$ | $\Delta\gamma$<br>$\Pr(SFRP5\vert MYC)$ | $\Delta\gamma$<br>$\Pr(MYC\vert SFRP5)$ |
| --- | --- | --- | --- | --- | --- |
| 0.8 | 0.1 | 0.125 | 0.117783 | -0.006463410 | 0.000000e+00 |
| 0.7 | 0.1 | 0.1428571 | 0.1335314 | -0.006967724 | 5.551115e-17 |
| 0.6 | 0.1 | 0.1666667 | 0.1541507 | -0.007515486 | -5.551115e-17 |
| 0.5 | 0.1 | 0.2 | 0.1823216 | -0.008112496 | 0.000000e+00 |
| 0.8 | 0.1 | 0.125 | 0.117783 | -0.029059115 | 0.000000e+00 |
| 0.7 | 0.2 | 0.2857143 | 0.2513144 | -0.022595705 | 0.000000e+00 |
| 0.6 | 0.3 | 0.5 | 0.4054651 | -0.015627982 | -5.551115e-17 |
| 0.5 | 0.4 | 0.8 | 0.5877867 | -0.008112496 | 0.000000e+00 |

**Figure 10:**
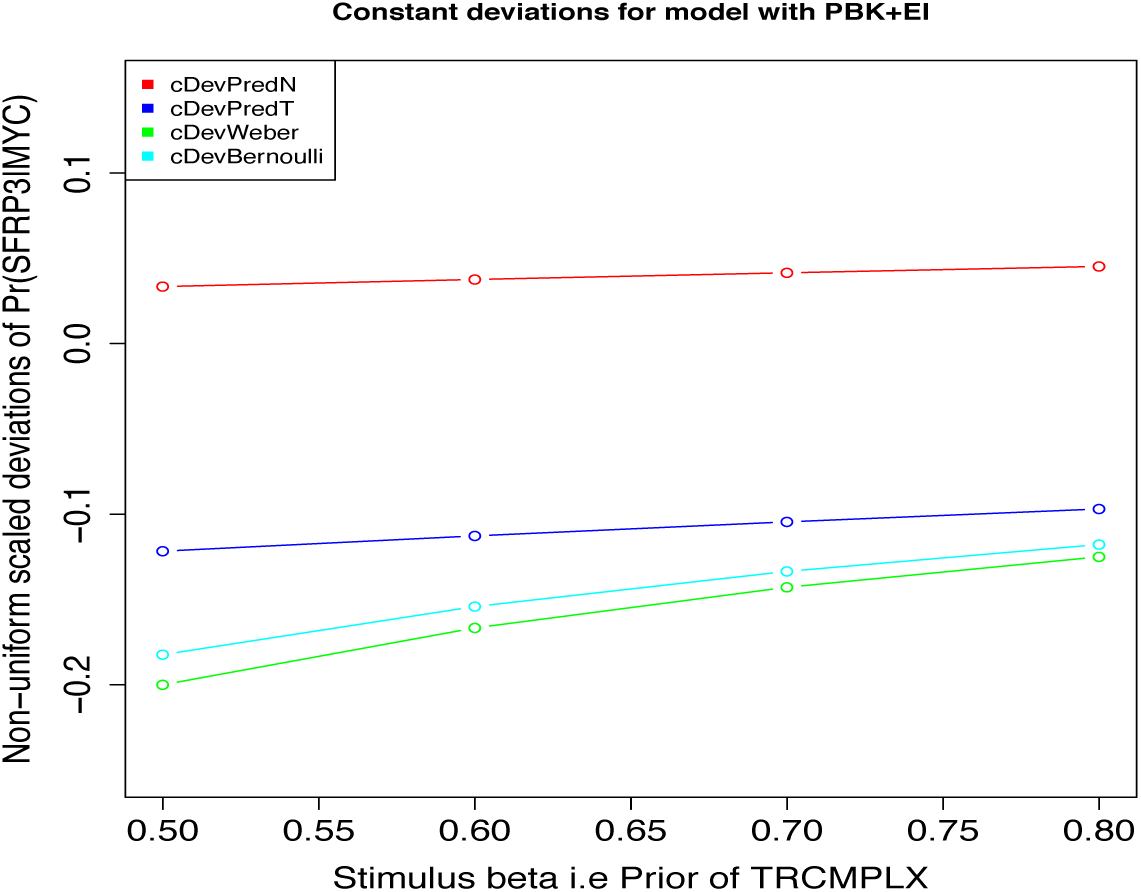
Constant deviations in *β* i.e ETGN and corresponding deviations in Pr(*SFRP* 3 | *MY C*) for both normal and tumor test samples. Corresponding Weber and Bernoulli deviations were also recorded. Note that the plots and the y-axis depict scaled deviations to visually analyse the observations. The model used is *M*_*PBK*+*EI*_. Red - deviation in Pr(*SFRP* 3 | *MY C*) in Normal case using Weber’s law, Blue - deviation in Pr(*SFRP* 3 | *MY C*) in Tumor using Weber’s law, Green - constant deviation in Webers law, Cyan - constant deviation in Bernoullis law.

Considering figure 10, when deviations are constant in both Weber and Bernoulli formulation, the deviations in the prediction of Pr(*SFRP* 3 | *MY C*) is observed to be logarithmic in the normal samples (apropos the Weber and Bernoulli deviations represented by green and cyan curves). Deviation in predictions are depicted by the red (blue) curves for normal (tumor) samples. Such a behaviour is not observed for Pr(*MY C* | *SFRP* 3) as is depicted in figure 11. Note that the interaction for *SFRP* 3 given *MY C* was observed to be reversible in normal and tumor cases. But this is not so with the interaction for *MY C* given *SFRP* 3. It might be expected that the non conformance of logarithmic-power law for Pr(*MY C* | *SFRP* 3) may be due to the non preservation/existence of the interaction of *MY C* given *SFRP* 3. This is so because Pr(*SFRP* 3 | *MY C*) depicts a reversible *SFRP* 3 *<>* −|*MY C* (*MY C <>* −| *SFRP* 3) in the network on normal (tumor) samples, while Pr(*MY C SFRP* 3) does not depict a reversible *MY C* | *<> SFRP* 3 (*MY C* | − | *SFRP* 3) in the network on normal (tumor) samples.

**Figure 11:**
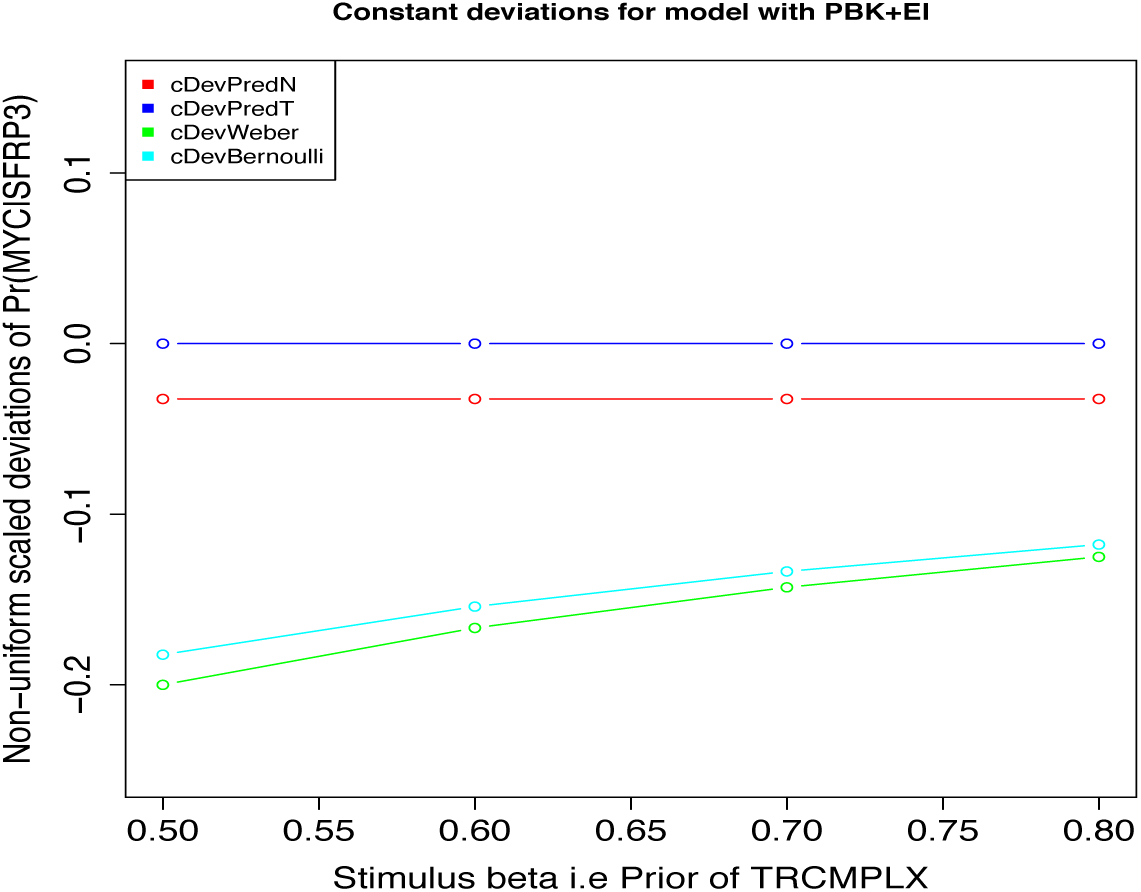
Same as figure 10 but for Pr(*MY C*|*SFRP*3).

Similar behaviour was observed in the case of incremental deviations as depicted in figures 12 and 13. Analysis of the behaviour of other gene-gene interactions showing *<>* −| or |− *<>* can be observed in a similar way and can be produced by executing the R code in Weber Fechner law.r provided in Google drive https://drive.google.com/folderview?id=0B7Kkv8wlhPU-T05wTTNodWNydjA&usp=sharing. Note that plots need manual axis and title adjustments. Some of the plot results has been compressed in the zip file titled Results-2015.zip.

**Figure 12:**
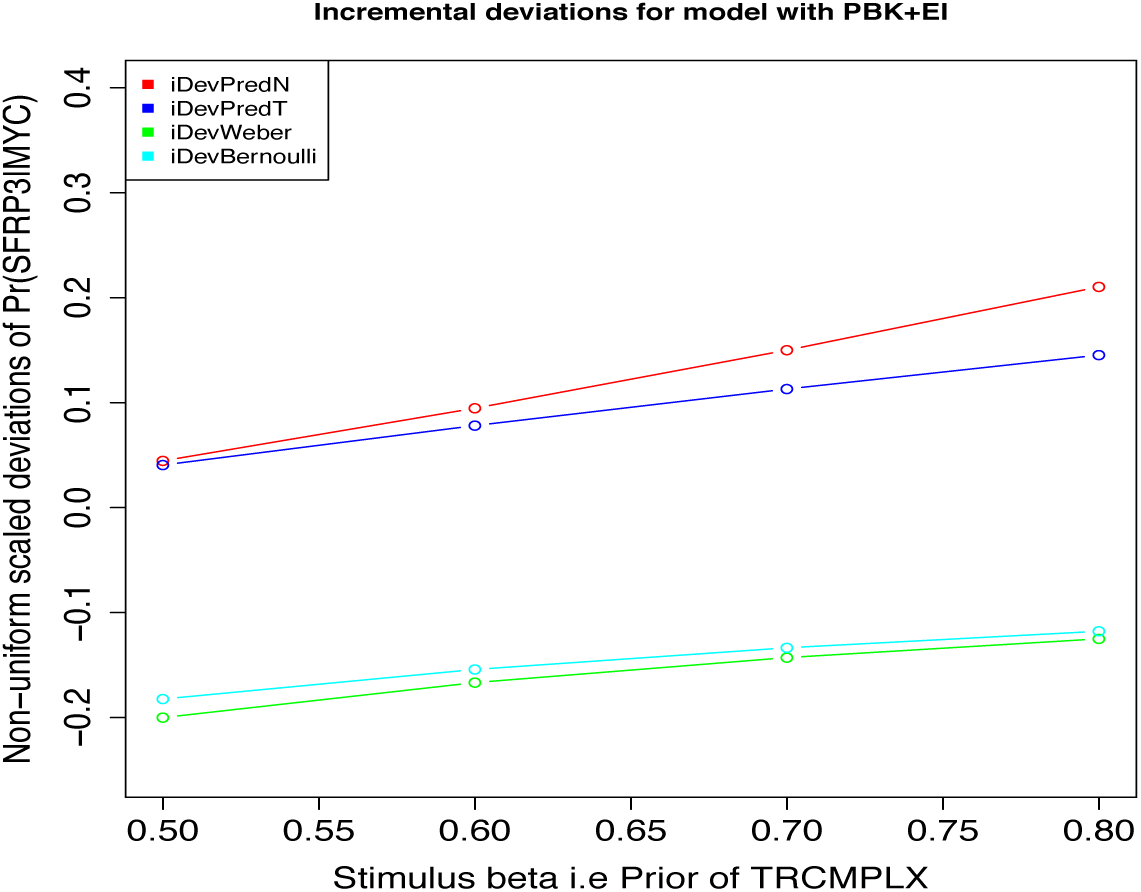
Same as figure 10 but for Pr(*SFRP* 3 *| MY C*). Instead of constant deviations, incremental deviations are represented.

**Figure 13:**
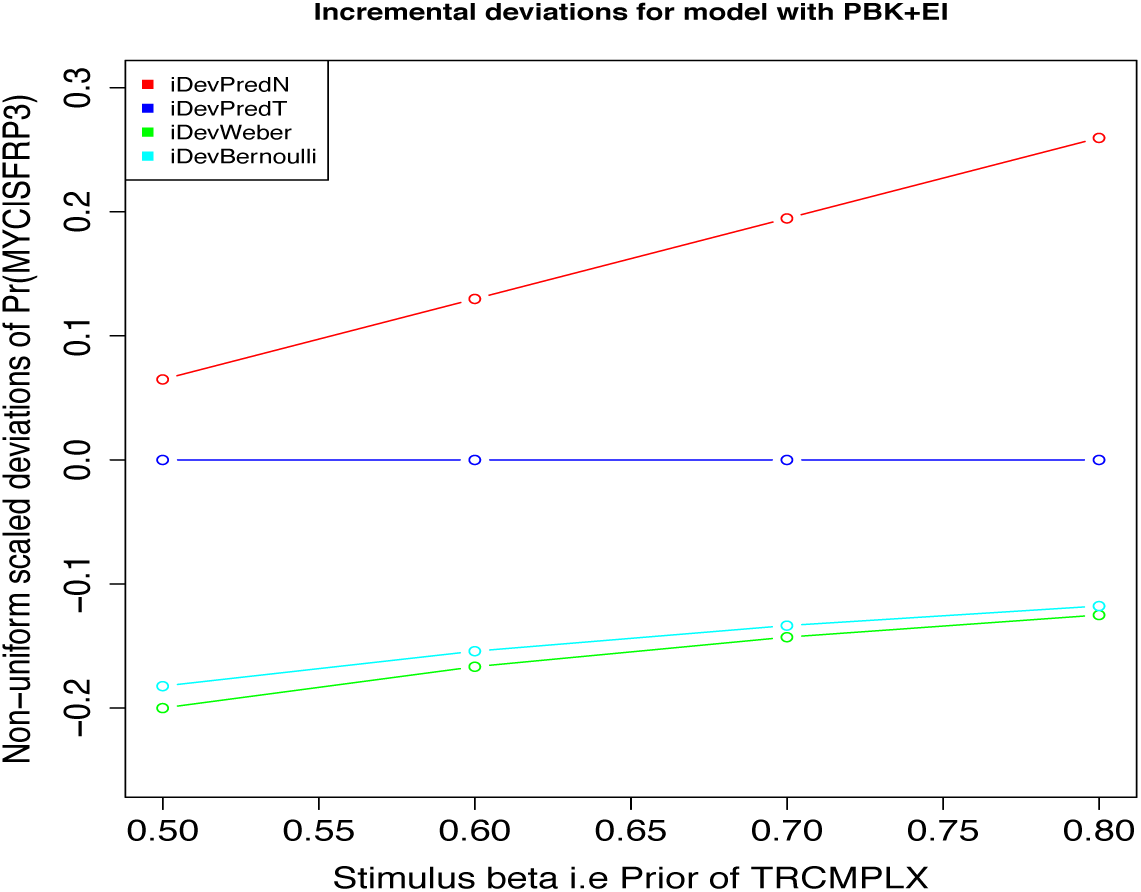
Same as figure 10 but for Pr(*MY C* | *SFRP* 3). Instead of constant deviations, incremental deviations are represented.

**Case:** | − | **or** *<>* − *<>* **with** *θ* = *θ*_*N*_

Again, as pointed out in Sinha (2014), the unknown behaviour of *SFRP* 2 in the Wnt pathway has been captured using computational causal inference. In figure 9, *SFRP* 2 shows preservation in the network and it’s interaction with other genetic factors involved in the model proposed in Sinha (2014) has been depicted. In one such paired interaction between *SFRP* 2 and *CD*44, both showed repression (activation) in normal (tumor) samples. As the change in the effect of transcription complex was induced via sensitizing the initially assigned cpt values, the deviations in the prediction of the gene-gene interaction network was observed to follow the logarithmic-power law crudely.

Table 9 and 10 show these deviations in the prediction of the interactions for both the normal and the tumor cases. The tables show how deviations are affected when the changes in the effect of the transcription complex are done at constant and incremental level. To summarize the results in these tables, graphs were plotted in figures 14 for Pr(*SFRP* 2 | *CD*44) (constant deviations), 15 for Pr(*CD*44 | *SFRP* 2) (constant deviations), 16 for Pr(*SFRP* 2 | *CD*44) (incremental deviations) and 15 for Pr(*CD*44 | *SFRP* 2) (incremental deviations).

**Table 9:** Deviation study for Pr(*SFRP* 2*|CD*44) and Pr(*CD*44*|SFRP* 2) for normal case

| $\beta$ | $\Delta\beta$ | $\frac{\Delta\beta}{\beta}$ | $\log(1 + \frac{\Delta\beta}{\beta})$ | $\Delta\gamma$<br>$\Pr(SFRP2\vert CD44)$ | $\Delta\gamma$<br>$\Pr(CD44\vert SFRP2)$ |
| --- | --- | --- | --- | --- | --- |
| 0.8 | 0.1 | 0.125 | 0.117783 | -0.0007505445 | 0.0002943409 |
| 0.7 | 0.1 | 0.1428571 | 0.1335314 | -0.0009398116 | 0.0002943409 |
| 0.6 | 0.1 | 0.1666667 | 0.1541507 | -0.0011360011 | 0.0002943409 |
| 0.5 | 0.1 | 0.2 | 0.1823216 | -0.0013397022 | 0.0002943409 |
| 0.8 | 0.1 | 0.125 | 0.117783 | -0.004166059 | 0.0011773636 |
| 0.7 | 0.2 | 0.2857143 | 0.2513144 | -0.003415515 | 0.0008830227 |
| 0.6 | 0.3 | 0.5 | 0.4054651 | -0.002475703 | 0.0005886818 |
| 0.5 | 0.4 | 0.8 | 0.5877867 | -0.001339702 | 0.0002943409 |

**Table 10:** Deviation study for Pr(*SFRP* 2*|CD*44) and Pr(*CD*44*|SFRP* 2) for tumor case

| $\beta$ | $\Delta\beta$ | $\frac{\Delta\beta}{\beta}$ | $\log(1 + \frac{\Delta\beta}{\beta})$ | $\Delta\gamma$<br>$\Pr(SFRP2\vert CD44)$ | $\Delta\gamma$<br>$\Pr(CD44\vert SFRP2)$ |
| --- | --- | --- | --- | --- | --- |
| 0.8 | 0.1 | 0.125 | 0.117783 | 0.02291329 | 0.01491512 |
| 0.7 | 0.1 | 0.1428571 | 0.1335314 | 0.02132802 | 0.01491512 |
| 0.6 | 0.1 | 0.1666667 | 0.1541507 | 0.01962443 | 0.01491512 |
| 0.5 | 0.1 | 0.2 | 0.1823216 | 0.01779600 | 0.01491512 |
| 0.8 | 0.1 | 0.125 | 0.117783 | 0.08166175 | 0.05966047 |
| 0.7 | 0.2 | 0.2857143 | 0.2513144 | 0.05874846 | 0.04474535 |
| 0.6 | 0.3 | 0.5 | 0.4054651 | 0.03742044 | 0.02983024 |
| 0.5 | 0.4 | 0.8 | 0.5877867 | 0.01779600 | 0.01491512 |

**Figure 14:**
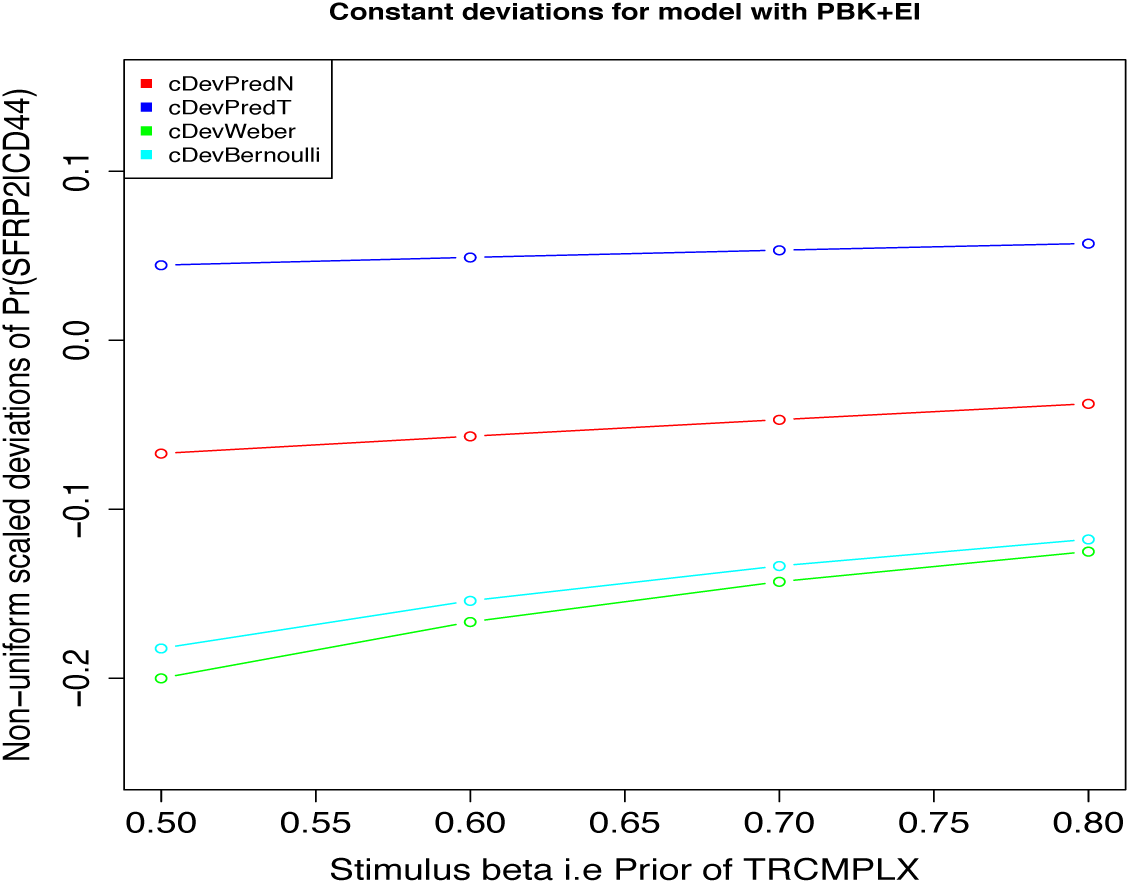
Constant deviations in *β* i.e ETGN and corresponding deviations in Pr(*SFRP* 2 | *CD*44) for both normal and tumor test samples. Corresponding Weber and Bernoulli deviations were also recorded. Note that the plots and the y-axis depict scaled deviations to visually analyse the observations. The model used is *M*_*PBK*+*EI*_. Red - deviation in Pr(*SFRP* 2 | *CD*44) in Normal case using Weber’s law, Blue - deviation in Pr(*SFRP* 2 | *CD*44) in Tumor using Weber’s law, Green - constant deviation in Webers law, Cyan - constant deviation in Bernoullis law.

Considering figure 14, when deviations are constant in both Weber and Bernoulli formulation, the deviations in the prediction of Pr(*SFRP* 2 | *CD*44) is observed to be logarithmic in the normal samples (apropos the Weber and Bernoulli deviations represented by green and cyan curves). Deviation in predictions are depicted by the red (blue) curves for normal (tumor) samples. Such a behaviour is not observed for Pr(*CD*44 | *SFRP* 2) as is depicted in figure 15. Even though Pr(*CD*44 | *SFRP* 2) was computationally estimated through a model, the interaction for *CD*44 given *SFRP* 2 was not observed in both normal and tumor cases while the interaction for *SFRP* 2 given *CD*44 was observed to be reversible. This points to a crucial fact that the interactions interpreted from conditional probabilities are not always two sided. Thus the interpretation for Pr(*g*_*i*_ | *g*_*j*_) is investigated in both directions as *g*_*i*_ *ℐ ℛ g*_*j*_ and *g*_*j*_ *ℐ ℛ g*_*i*_ to get a full picture. Not that the results are wrong, but all angles of interpretations need to be investigated to get the picture between any two genes. Similar behaviour was observed in the case of incremental deviations as depicted in figures 16 and 17. Note that graph for incremental deviation in Pr(*CD*44 *| SFRP* 2) is just a cumulative effect and does not state anything about the logarithmic law.

**Figure 15:**
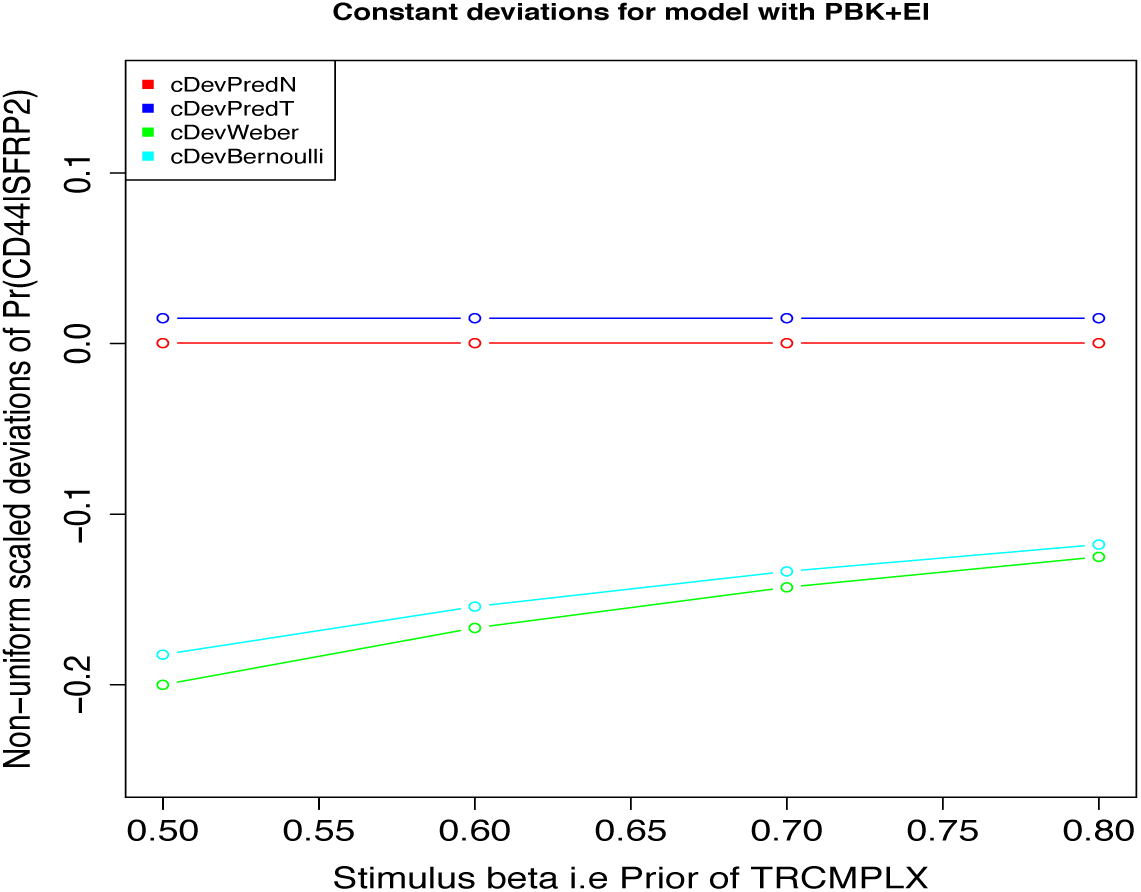
Same as figure 14 but for Pr(*CD*44*|SFRP* 2).

**Figure 16:**
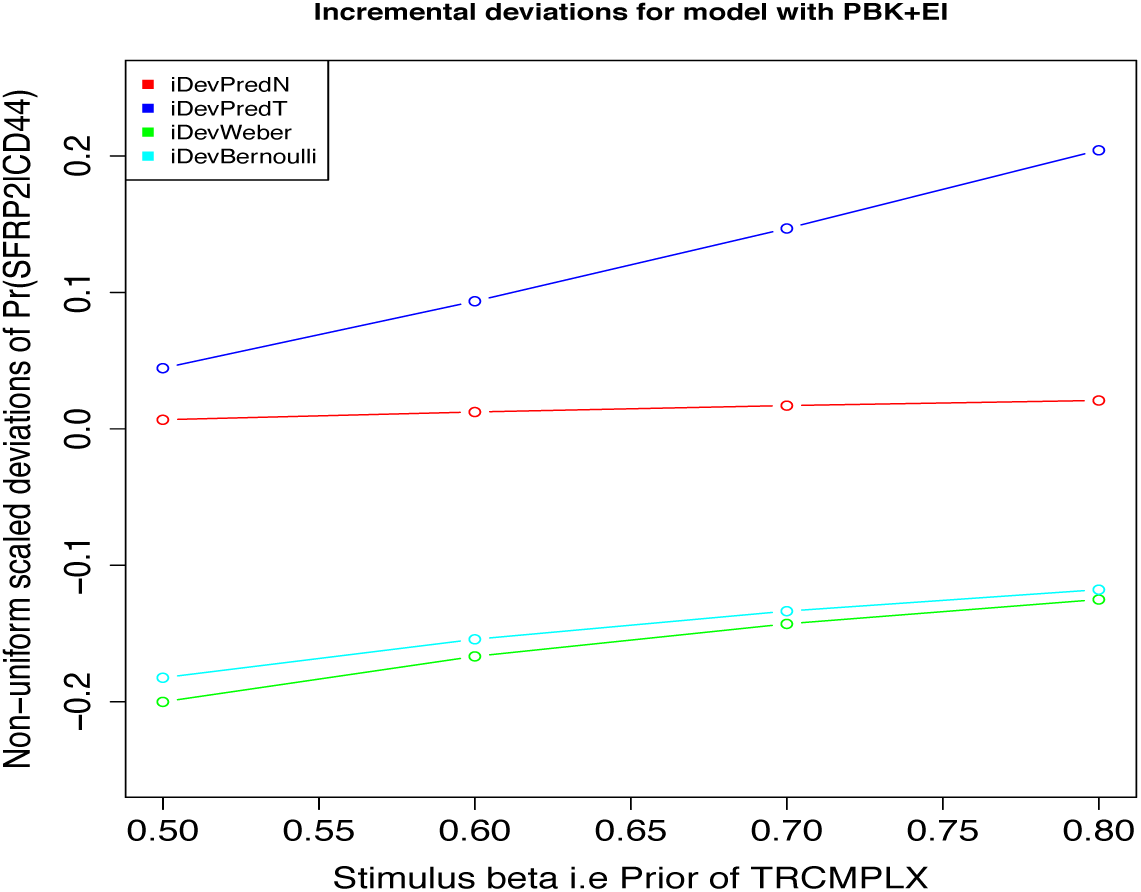
Same as figure 14 but for Pr(*SFRP* 2 | *CD*44). Instead of constant deviations, incremental deviations are represented.

**Figure 17:**
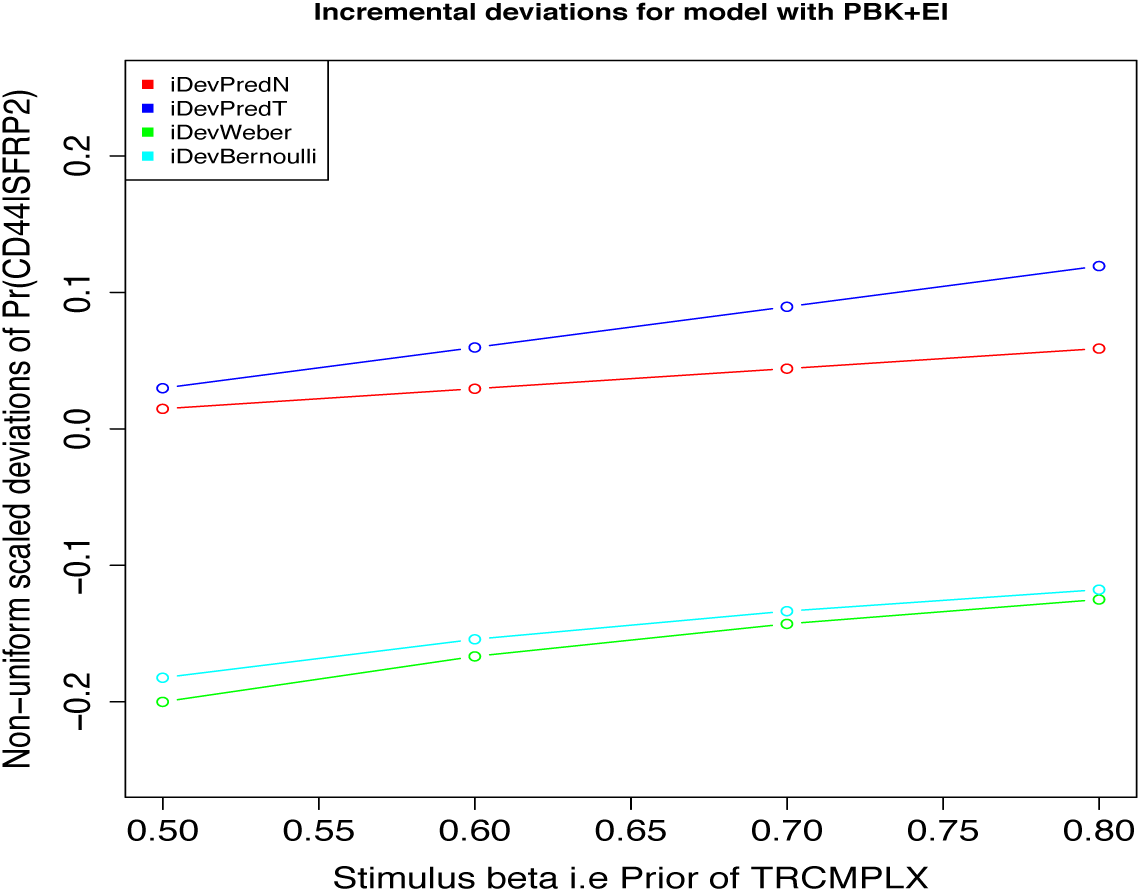
Same as figure 15 but for Pr(*CD*44 | *SFRP* 2). Instead of constant deviations, incremental deviations are represented.

Finally, note that the predicted conditional probability a gene *i* given evidence for gene *j* does not change but the inferred gene-gene interactions do change depending on the choice of the threshold. These changes are depicted in the figures 8 (table 4)and 9 (table 5). Dual interactions were inferred using the weighted mean as a discretization factor, as is shown next. These are dual interactions are marked in red colour in figure 9.

**Case: Dual interactions with** *θ* = *θ*_*N*_

The dual interactions revealed using weighted means indicate an important phenomena between any two genes. These interactions reveal that gene activation interplay might not always be constant for normal (tumour) samples. These in silico observations imply that a gene that was found to be actively expressed in normal sample might reverse activity at some stage or the other (an vice versa). Here, one such interaction is discussed in detail. Interpretations of the other dual interactions can be done in the same way. Results for other interactions are available but not presented here.

Also, a point to be observed is that the weighted means show much more crisp discretization during inference of gene-gene interaction in comparison to use of an arbitrary value of 0.5. To determine this distinction between the inferred gene-gene interactions obtained via weighted threshold and the arbitrary threshold of 0.5, the receiver operator curves (ROC) along with its corresponding area under the curve (AUC) are plotted. The ROCs are plotted using the discretized predicted values and the discretized labels obtained using the thresholds (computed from the training data) on the test data. The ROC graphs and their respective AUC values indicate how the predictions on the test data behaved under different values assigned to the TRCMPLX while training. Ideally, high values of AUC and steepness in ROC curve indicate good quality results. Finally, two sample Kolmogorov-Smirnov (KS) test was employed to measure the statistical significance between the distribution of predictions. If the cumulative distributions are not similar the KS test returns a small p-value. This small p-value indicates the existing statistical significance between the distributions under consideration.

Finally the ROC plots and AUC values for dual gene-gene interactions are also plotted and KS test is conducted to find the existence of statistical significance if any. These reveal the significance of existence of dual interactions in the signaling pathway which might not have been revealed using the arbitrary threshold value of 0.5. Plots are made using functions from the PRROC package provided by Grau et al. (2015).

**Interaction between DKK1 and DACT2 using** *θ* ∈ {*θ*_*N*_*, θ*_*T*_}Dual interactions *DACT* 2 *<> <> DKK*1 and *DKK*1 | − *<> DACT* 2 (*DACT* 2 | − *DKK*1 and *DKK*1 *<>* − | *DACT* 2) in normal (tumor) sample were found as depicted in figure 9. Figure 18 shows the kernel density estimate of the predicted conditional probabilities for both normal and tumor test cases. Using the weighted mean of the discretized values of the test samples (discretization done using median estimated from the training data as mentioned before), the predicted Pr(*DKK*1*|DACT* 2) and Pr(*DACT* 1*|DKK*1) are classified as active or passive. It might be useful to note that instead of using 0.5 as an arbitrary value, the weighted mean captures the distribution of labels in a much more realistic manner and helps infer interactions among the factors in the Wnt pathway.

**Figure 18:**
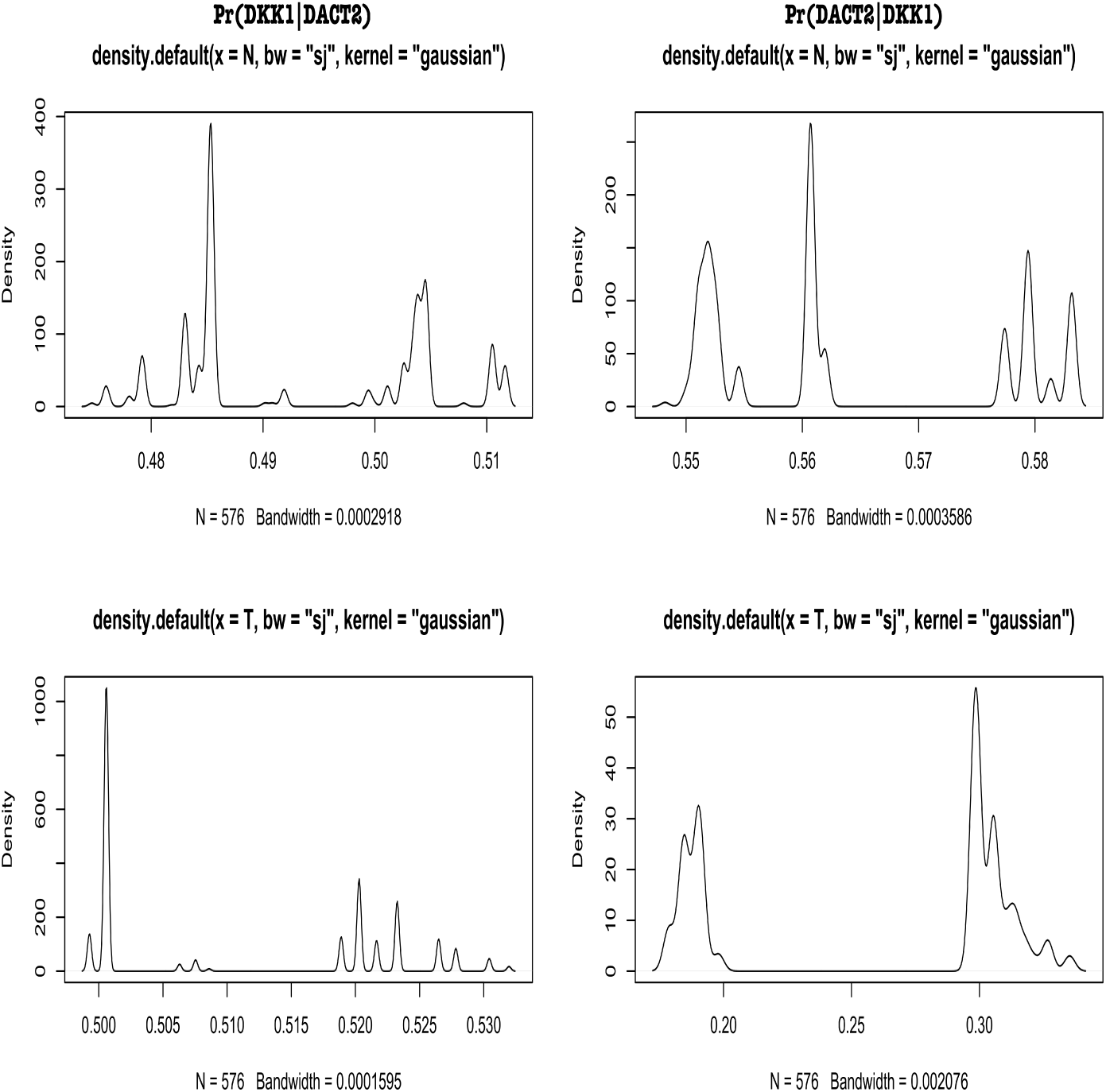
Kernel density estimates for predicted Pr(*DKK*1 | *DACT* 2) and Pr(*DACT* 2 | *DKK*1) in Normal and Tumor cases. Gaussian kernel is used for smoothing the density estimate. The bandwidth of the kernel is selected using the pilot estimation of derivative as proposed by Sheather and Jones (1991) and implemented in *R* programming language.

Note the distributions depicted in figure 18. In the first column of the figure, the median for Pr(*DKK*1 | *DACT* 2) in normal (tumor) case is 0.4853088 (0.5006437). These medians point to the mid value of the belief in the genegene interaction depicted by the range of predicted conditional probabilities. The weighted threshold 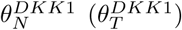 based on labels for normal (tumor) test case was estimated at 0.5138889 (0.4861111). The estimations come from the following computations in Equation 9 -

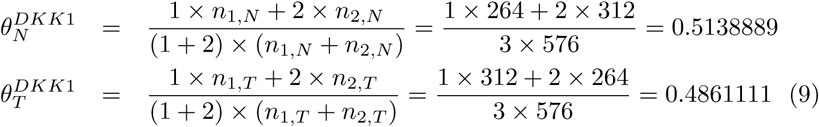

Similarly, in the second column of the figure, the median for Pr(*DACT* 2 | *DKK*1) in normal (tumor) case is 0.5606946 (0.2985911). The weighted threshold 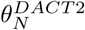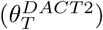 based on labels for normal (tumor) test case was estimated at 0.4583333 (0.5416667). The estimations come from the following computations in Equation 10 -

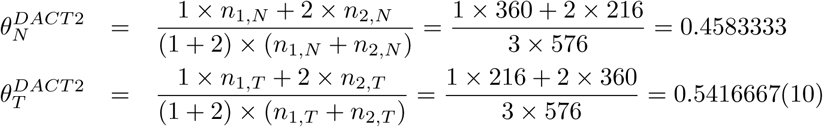

It can be observed that the discretization is more realistic and strict using the weighted threshold rather than using the arbitrary value of 0.5. The multiple peaks point to the different frequencies at which the predicted probabilities were recorded. Note that the probabilities here represent the belief in the activation status and the discretization only calibrates the belief into active and repressed state. To evaluate the results further wet lab tests are needed.

Using these distributions and distributions obtained using arbitrary value, the respective ROC are plotted and corresponding AUC values estimated. Finally, KS test is used to find the existence of statistical significance between the valid permutations of the distributions. These estimates further help derive insights about the interactions at a computational level. Figure 19 shows the ROC plots and the respective AUC values for the dual interactions observed via the in silico experiments. The following are compared -

1. labels of test data *ge*_*N*_ and discretized values of Pr(*DKK*1*|DACT* 2) and Pr(*DACT* 2*|DKK*1) using weighted mean in Normal case
2. labels of test data *ge*_*T*_ and discretized values of Pr(*DKK*1*|DACT* 2) and Pr(*DACT* 2*|DKK*1) using weighted mean in Tumor case

**Figure 19:**
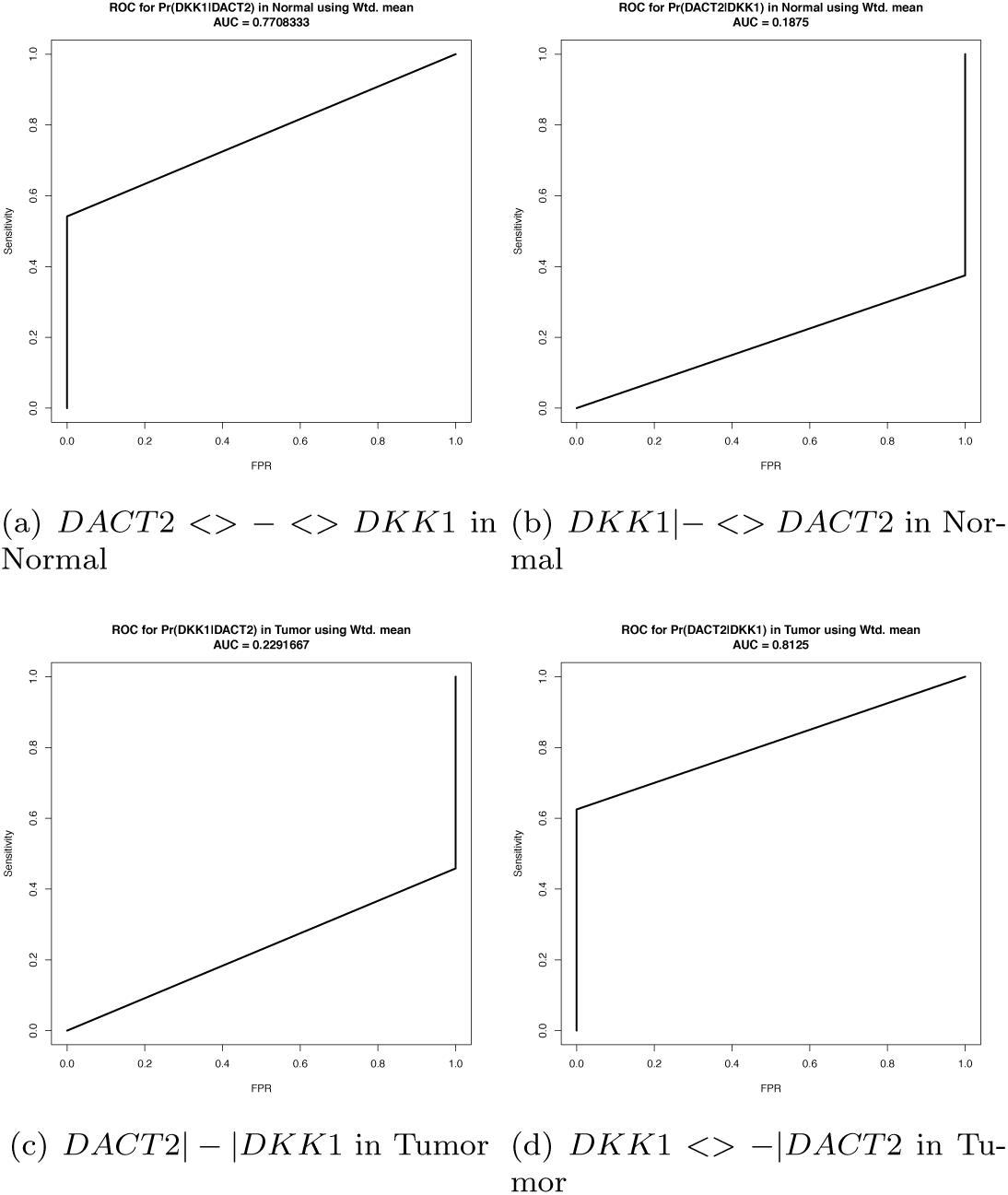
Column wise ROCs for Pr(*DKK*1 | *DACT* 2) (1^*st*^ column) and Pr(*DACT* 2 | *DKK*1) (2^*nd*^ column) have been plotted with ETGN value for the 90%. Row wise the plots depict the curves generated using weighted mean for Normal case and weighted mean for Tumor case. Respective AUC values for the ROC curves appear on the title of each of the graphs.

In figure 19 column wise the ROCs for Pr(*DKK*1 | *DACT* 2) (1^*st*^ column) and Pr(*DACT* 2 | *DKK*1) (2^*nd*^ column) have been plotted with ETGN value for the 90%. Row wise the plots depict the curves generated using weighted mean for Normal case and weighted mean for Tumor case. It can be seen that using the weighted mean, the subfigure 19(a) and 19(d) convey a good guess regarding the type of interaction prevailing in normal and tumor case. Thus *DACT* 2 *<> − <> DKK*1 i.e Pr(*DKK*1 | *DACT* 2) is highly favoured in Normal case while *DKK*1 *<>* −| *DACT* 2 i.e Pr(*DACT* 2 | *DKK*1) is highly favoured in Tumor case. Why this is so is because the normal cases show better results in terms of prediction in comparison to the tumor cases. This points to the fact that the interaction *DACT* 2 *<>* − *<> DKK*1 is strongly supported in the normal case in comparison to *DACT* 2 | − | *DKK*1 which is weakly supported in the tumor case. Even though the algorithm showed that interaction was reversible at computational level, ROC curves and corresponding AUC values indicate weakness in the belief that *DACT* 2 | − *| DKK*1 prevails in tumor cases. On the other hand, the interaction depicted by Pr(*DACT* 2 | *DKK*1) shows higher predictive quality in the tumor case with respect to the normal case. This means that *DKK*1 *<>* − *| DACT* 2 has more weight in tumor case than its reversible *DKK*1 | − *<> DACT* 2 counter part in the normal case. Taken together, the dual interactions do exist but with different strengths of belief as shown conditional probability values. The curves in sub-figure 19(b) and 19(c) indicate a bad guess and thus do not support the interactions *DKK*1| − *| <> DACT* 2 i.e Pr(*DACT* 2 | *DKK*1) in Normal case and *DACT* 2 | − *| DKK*1 i.e Pr(*DKK*1 | *DACT* 2) in Tumor case.

**Interaction between DKK1 and DACT2 using** *θ* = 0.5

In comparison to use of the weighted *θ*, the analysis of single interaction using *θ* = 0.5 is also presented. Figure 8 shows the interaction between *DKK*1 and *DACT* 2 as *DACT* 2 *<>* −| *DKK*1, i.e Pr(*DKK*1 | *DACT* 2). Using a 0.5 threshold on 18 it is possible to see that discretization of kernel density estimates of Pr(*DKK*1 | *DACT* 2) induces a degree of belief which is not exactly 0(1). This is not the case with Pr(*DACT* 2 | *DKK*1), were the discretization leads to an exact 0(1) which removes the degree of belief. Bayesian networks often represent the degree of belief in terms of some real valued number and exact probabilities of 0(1) are considered with suspicion.

Figure 20 shows the ROC plots and the respective AUC values for the dual interactions observed via the in silico experiments. The following are compared -

1. labels of test data *ge*_*N*_ and discretized values of Pr(*DKK*1*|DACT* 2) and Pr(*DACT* 2*|DKK*1) using arbitrary value of 0.5 in Normal case
2. labels of test data *ge*_*T*_ and discretized values of Pr(*DKK*1*|DACT* 2) and Pr(*DACT* 2*|DKK*1) using arbitrary value of 0.5 in Tumor case

**Figure 20:**
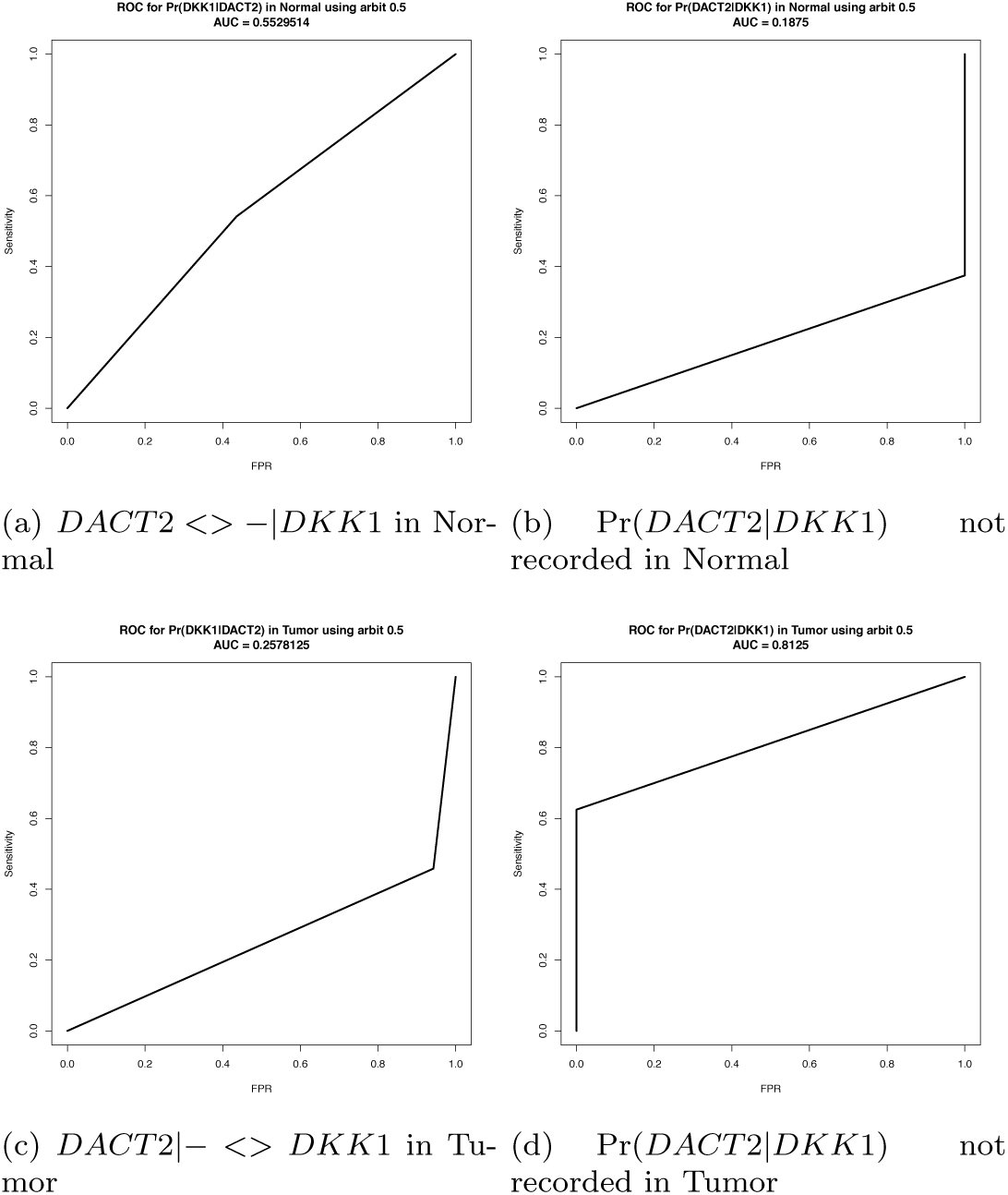
Column wise ROCs for Pr(*DKK*1 *| DACT* 2) (1^*st*^ column) and Pr(*DACT* 2 *| DKK*1) (2^*nd*^ column) have been plotted with ETGN value for the 90%. Row wise the plots depict the curves generated using arbit value of 0.5 for Normal case and arbit value of 0.5 for Tumor case. Respective AUC values for the ROC curves appear on the title of each of the graphs.

In figure 20 column wise the ROCs for Pr(*DKK*1 | *DACT* 2) (1^*st*^ column) and Pr(*DACT* 2 | *DKK*1) (2^*nd*^ column) have been plotted with ETGN value for the 90%. Row wise the plots depict the curves generated using arbitrary threshold of 0.5 for Normal case and Tumor case. It can be seen that using the a value of 0.5, the subfigure 20(a) conveys a negligibly good guess regarding the type of interaction prevailing in normal. Thus *DACT* 2 *<>* − | *DKK*1 i.e Pr(*DKK*1 | *DACT* 2) is highly favoured in Normal case. On the other hand the 20(c) conveys a very bad guess regarding the reversal of interaction in Tumor case for Pr(*DKK*1 | *DACT* 2). Finally, it was noted that the degree of belief in Pr(*DACT* 2 | *DKK*1) was not at all recorded via thresholding. Thus even though 20(b) and 20(d) show recorded ROCs but the discretization of 0.5 does not capture the involved interaction. Thus the arbitrary value of 0.5 is not a good factor for inferring interactions.

Comparing figures 20 and 19, it is clear that the later gives a better guess in terms of the interpretation of the interaction obtained by discretizing the kernel density estimates of inferred conditional probabilities. To evaluate the statistical significance of the predicted probabilities, the values of the KS test are tabulated and analyzed. Table 11 represents the computed values. The first four rows show the existing significance between the predictions for which the ROC curves have be plotted and described earlier. The next describes the significance between predictions based on thresholds for both normal and tumor cases. Note that some tests show no significance at all as is the case with Pr(*DACT* 2 *| DKK*1). In general, significance values differ depending on different interactions. Finally, significance values between interactions are also tabulated. It was found that there exists statistical difference between the inferred dual interactions as shown by the low p-values. Similar interpretations can be derived and respective measures can be plotted from the in silico observations.

**Table 11:** Kolmogorov-Smirnov test indicating statistical significance between the distribution of predictions. Statistical significance is evaluated by observing the p-value. Small p-value indicates that significant difference. Significance test is conducted between (1) discretized values of predictions and existing test labels (2) discretized values of predictions based on weighted threshold and discretized values of predictions based on arbit threshold and (3) between predictions representing the dual interactions (obtained using both thresholds). *I*_1_ and *I*_2_ correspond to interactions inferred from Pr(*DKK*1 *| DACT* 2) and Pr(*DACT* 2 *| DKK*1), respectively.

| Kolmogorov-Smirnov test |  |  |  |  |
| --- | --- | --- | --- | --- |
| Sr. No. | Discretized Val. vs Labels | p-value | Discretized Val. vs Labels | p-value |
| | $\Pr(DKK1\vert DACT2)$ | | $\Pr(DACT2\vert DKK1)$ | |
| 1. | wtd. mean (N) vs $ge_N$ | D = 0.5417<br>p-value < $2.2e^{-16}$ | wtd. mean (N) vs $ge_N$ | D = 0.625<br>p-value < $2.2e^{-16}$ |
| 2. | wtd. mean (N) vs $ge_N$ | D = 0.1059<br>p-value = 0.003129 | wtd. mean (N) vs $ge_N$ | D = 0.625<br>p-value < $2.2e^{-16}$ |
| 3. | wtd. mean (N) vs $ge_T$ | D = 0.5417<br>p-value < $2.2e^{-16}$ | wtd. mean (T) vs $ge_T$ | D = 0.625<br>p-value < $2.2e^{-16}$ |
| 4. | wtd. mean (N) vs $ge_T$ | D = 0.4844<br>p-value < $2.2e^{-16}$ | wtd. mean (T) vs $ge_T$ | D = 0.625<br>p-value < $2.2e^{-16}$ |
| KS test between predictions using wtd. mean and arbitrary value of 0.5 |  |  |  |  |
| Sr. No. | $\Pr(DKK1\vert DACT2)$ | KS value | $\Pr(DKK2\vert DACT1)$ | KS value |
| 1. | wtd. mean vs arbit. (N) | D = 0.4358<br>p-value < $2.2e^{-16}$ | wtd. mean vs arbit. (N) | D = 0<br>p-value = 1 |
| 2. | wtd. mean vs arbit. (T) | D = 0.0573<br>p-value = 0.3009 | wtd. mean vs arbit. (T) | D = 0<br>p-value = 1 |
| KS test between predictions of interactions $I_1$ and $I_2$ | | | | |
| 1. | wtd. mean - $I_1$ (N) vs $I_2$ (N) | D = 1<br>p-value < $2.2e^{-16}$ | wtd. mean - $I_1$ (T) vs $I_2$ (T) | D = 1<br>p-value < $2.2e^{-16}$ |
| 2. | arbit. - $I_1$ (N) vs $I_2$ (N) | D = 0.5642<br>p-value < $2.2e^{-16}$ | arbit. - $I_1$ (T) vs $I_2$ (T) | D = 0.9427<br>p-value < $2.2e^{-16}$ |

## 7. Caveats

This work does not take into account the time series data which contains much more crucial information rather than the static data of gene expression. The inferences have been made regarding a natural phenomena based on the exploration of a computational causal model via sensitivity analysis. The results discussed are based on deviations of inferred conditional probabilities which encode a degree of belief in the occurrence of an event. Even if dynamic bayesian models are used, the observations will be made on degree of beliefs only. Also, the current bayesian network model does not encode the cyclic feedback loops. This has serious implications in the fact that the model might not capture correct interactions. The problem can be overcome to a certain extent by encoding the biological knowledge such that concepts of *d*-connectivity/separability exploit the inherent prior knowledge and thus help in proper inferences. More specifically, the model captures a snapshot in time but by varying the parameters or the prior/conditional probability tables, it is possible to verify the natural phenomena under investigation.

## 8. Future directions

In context of the above observations, dynamic models might reveal greater information regarding the psychophysical laws. Work by Goentoro and Kirschner (2009) employs sensitivity analysis methods to reveal such laws by tuning single parameters. There might be a few ways to measure fold change in single an multi parameter settings. Future work might involve deeper study of the phenomena based on multi-parameter setting in a dynamic bayesian network model. If one incorporates nodes in between two time snapshots of *β*-*catenin* concentration in a dynamic bayesian network, one might be able to measure the changes at different phases of the signaling pathway. For example, in figure 21 a set of nodes measuring the different concentrations of *β*-*catenin* (say *N*) are depicted. In a dynamic bayesian network, the previous concentration at *t* is connected to the next concentration at *t* + 1. Also, to measure the effect of difference (Δ*N*), a change in concentration can be measured. Computations regarding fold change (Δ*N*) could then be estimated as posterior probabilities given the two concentrations, which the Bayesian networks can easily handle. In case more parameters need to be involved (say the effect of Wnt and APC together), nodes might be added as shown below. Then the fold change is conditional on *N* (*t* + 1), *N* (*t* + 2), Δ*Wnt* and Δ*APC* and is estimated as Pr(Δ*N* (*t* + 1) *N* (*t* + 1)*,N* (*t* + 2), Δ*Wnt,* Δ*APC*).

**Figure 21:**
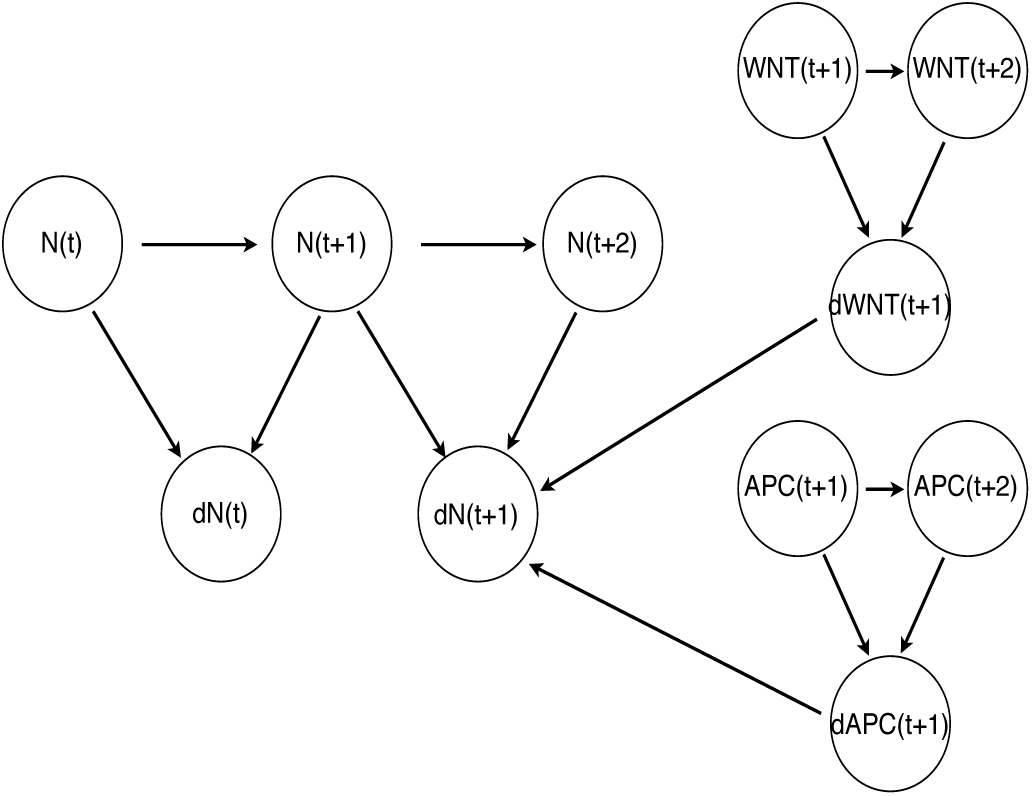
A schematic diagram of a dynamic bayesian network model that might help study the fold change and the logarithmic psychophysical laws behind the changes.

Regarding sensitivity analysis, in nonlinear problems, it might be useful to use Sobol’ (1990) indices to estimate the sensitivity of the parameters. These indices are a way to estimate the changes in a multiparameter setting thus helping one to conduct global sensitivity analysis instead of local sensitivity analysis Glen and Isaacs (2012). Finally, with respect to the robustness of the gene-gene interaction network, the current work employs a very simple algorithm to construct the network and infer preserved interactions across the range of values set for a particular parameter. This helps in eliminating interactions that do not contribute enough biological information in the pathway or are non existant and require further analysis by integration of more data. Work in these lines would require incorporation of bigger datasets.

## 9. Conclusions

In this preliminary work via sensitivity analysis, the variation in predictive behaviour of *β*-*catenin* based transcription complex conditional on gene evidences is shown to follow power-logarithmic psychophysical law crudely. This implies deviations in output are proportional to increasing function of deviations in the input and show constancy for higher values of input. This points towards stability in the behaviour of transcriptional activity downstream of the Wnt pathway. As a further development, computational analysis shows that the preserved gene-gene interactions are also subject to these power-logarithmic psychophysical laws. The prevalence of these laws is reported for interaction between elements of pairs of (*SFRP* 3, *MY C*), (*SFRP* 2, *CD*44) and (*DKK*1, *DACT* 2). As a precursor to the analysis of these laws at interaction level, the biologically inspired epigenetically influenced computational causal models were used to infer gene-gene interaction from conditional probabilities of individual gene activation given the status of another gene activation. In relation of colorectal cancer cases, it is now possible to infer the type of interaction that might be happening among the genes at a pair wise level using BN models and further wet lab studies can be developed to investigate the inferred prevalence of power-logarithmic psychophysical laws at interaction level within the pathway. To assert the fact, in a recent development via wet lab experiments by Olsman and Goentoro (2016), it has been confirmed that there are existence of sensors that behave in a logarithmic fashion. The wet lab work by Olsman and Goentoro (2016) supports the earlier proposed crude postulates based on computational sensitivity analysis of this manuscript regarding the existence of logarithmic behaviour in the signaling pathways. It also signifies the importance of systems biology approach where in silico experiments combined with in vivo/in vitro experiments have the power to explore the deeper mechanisms of a signaling pathway.

## Acknowledgement

Thanks to - • Dr. Uri Alon from Weizmann Institute of Science, Israel, for lively and critical discussion on behaviour of curves at the International Conference on Systems Biology of Human Disease (2015), held at German Cancer Research Center in Heidelberg, Germany. • Dr. Silja Renooij for providing useful feedback and editing the manuscript. • the Royal Society of Chemistry (RSC) for giving permission to reproduce parts of material in S. Sinha, Integr. Biol., 2014, DOI: 10.1039/C4IB00124A. • all anonymous reviewers who have helped in refining this manuscript.

## Declaration of Interest

No conflict of interest.

## Appendix

### 9.1. Steps for construction of gene gene interaction networks

Before starting the construction of interactions from the conditional probabilities, assign a variable *gg*_*I*_ as an empty list (say in R language). Then *∀i* genes, execute the following -

1. ∀ 576 runs iterated by a counter *j*
  a. append *x*_*N*_ with the vector whose elements are Pr(*g*_*i*_ = active | *g*_*k*_ evidence) *∀k* genes in the *j*^*th*^ run for Normal test sample. This creates a matrix at the end of the runs.
  b. append *x*_*T*_ with the vector whose elements are Pr(*g*_*i*_ = active | *g*_*k*_ evidence) *∀k* genes in the *j*^*th*^ run for Tumor test sample. This creates a matrix at the end of the runs.
  c. append *ge*_*N*_ with the vector whose elements are *ge*_*k*_ evidence ∀*k* genes in the *j*^*th*^ run for Normal test sample. This creates a matrix at the end of the runs.
  d. append *ge*_*T*_ with the vector whose elements are *ge*_*k*_ evidence ∀*k* genes in the *j*^*th*^ run for Tumor test sample. This creates a matrix at the end of the runs.
2. assign variables *ge*, *aa*_*N*_, *ar*_*N*_, *ra*_*N*_, *rr*_*N*_, *aa*_*T*_, *ar*_*T*_, *ra*_*T*_, *rr*_*T*_, *PggN*, *PggT* to an empty vector *c*() (say in R language). Note - a (r) means activation (repression).
3. compute mean across columns of *xN* and *xT* to obtain averaged 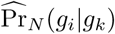and 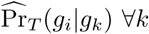 gene evidences and *i* genes. Note *k, i* ∈ 1*, …, n* if *n* is the total number of genes.
4. assign a vector of 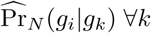 *∀k* genes to *PggN* and a vector of 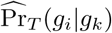 genes to *PggT*
5. *∀k* genes except the *i*^*th*^ one
  a. if(*k* ≠ *i*)
    i. assign variables *tmpaa*_*N*_, *tmpar*_*N*_, *tmpra*_*N*_, *tmprr*_*N*_, *tmpaa*_*T*_, *tmpar*_*T*_, *tmpra*_*T*_ and *tmprr*_*T*_ to 0.
    ii. assign threshold values *θ* to either a fixed value (say 0.5) or a weighted mean.
    iii. if assigning a weighted mean, compute the threshold *θ*_*N*_ as the weighted mean of the labels of the test data i.e evidences for the *i*^*th*^ gene, in the case of Normal samples (top formula in Equation 8). Similarly, compute the threshold *θ*_*T*_ as the weighted mean of the labels of the test data i.e evidences for the *i*^*th*^ gene, in the case of Tumor samples (bottom formula in Equation 8).
    iv. *∀* 576 runs iterated by a counter *l*
      A. if(*ge*_*N*_[*l*,*k*] == 1 and *x*_*N*_[*l*,*k*] *< θ*) increment *tmprr*_*N*_ by 1
      B. else if(*ge*_*N*_[*l*,*k*] == 1 and *x*_*N*_[*l*,*k*] *>*= *θ*) increment *tmpar*_*N*_ by 1
      C. else if(*ge*_*N*_[*l*,*k*] == 2 and *x*_*N*_[*l*,*k*] *< θ*) increment *tmpra*_*N*_ by 1
      D. else if(*ge*_*N*_[*l*,*k*] == 2 and *x*_*N*_[*l*,*k*] *>*= *θ*) increment *tmpaa*_*N*_ by 1
      E. if(*ge*_*T*_ [*l*,*k*] == 1 and *x*_*T*_ [*l*,*k*] *< θ*) increment *tmprr*_*T*_ by 1
      F. else if(*ge*_*T*_ [*l*,*k*] == 1 and *x*_*T*_ [*l*,*k*] *>*= *θ*) increment *tmpar*_*T*_ by 1
      G. else if(*ge*_*T*_ [*l*,*k*] == 2 and *x*_*T*_ [*l*,*k*] *< θ*) increment *tmpra*_*T*_ by 1
      H. else if(*ge*_*T*_ [*l*,*k*] == 2 and *x*_*T*_ [*l*,*k*] *>*= *θ*) increment *tmpaa*_*T*_by 1
    v. Comment - store results
    vi. append *ge* with *g*_*k*_, *rr*_*N*_ with *tmprr*_*N*_, *ar*_*N*_ with *tmpar*_*N*_, *ra*_*N*_ with *tmpra*_*N*_, *aa*_*N*_ with *tmpaa*_*N*_, *rr*_*T*_ with *tmprr*_*T*_, *ar*_*T*_ with *tmpar*_*T*_, *ra*_*T*_ with *tmpra*_*T*_ and *aa*_*T*_ with *tmpaa*_*T*_
  b. store the variables in the previous step to a data frame (say in R language) to a variable *stats*.
  c. Comment - 1 means aa, 2 means ar, 3 means ra, 4 means rr
  d. assign variables *gg*_*IN*_ and *gg*_*IT*_ as empty vector []
  e. *∀j* gene except the *i*^*th*^ one under consideration
    i. find the index *idx*_*N*_ in stats that corresponds to 1 or 2 or 3 or 4
    ii. if(*idx*_*N*_ == 1) append *gg*_*IN*_ with interaction string *stats§g*_*j*_ *<>* − *<> g*_*i*_
    iii. else if(*idx*_*N*_ == 2) append *gg*_*IN*_ with interaction string *stats§ge*_*j*_*|*− *<> g*_*i*_
    iv. else if(*idx*_*N*_ == 3) append *gg*_*IN*_ with interaction string *stats§g*_*j*_ *<>* −*|g*_*i*_
    v. else if(*idx*_*N*_ == 4) append *gg*_*IN*_ with interaction string *stats§g*_*j*_*|*−*|g*_*i*_
    vi. find the index *idx*_*N*_ in stats that corresponds to 1 or 2 or 3 or 4
    vii. if(*idx*_*T*_ == 1) append *gg*_*IT*_ with interaction string *stats§g*_*j*_ *<>* − *<> g*_*i*_
    viii. else if(*idx*_*T*_ == 2) append *gg*_*IT*_ with interaction string *stats§g*_*j*_*|*− *<> g*_*i*_
    ix. else if(*idx*_*T*_ == 3) append *gg*_*IT*_ with interaction string *stats§g*_*j*_ *<>* −*|g*_*i*_
    x. else if(*idx*_*T*_ == 4) append *gg*_*IT*_ with interaction string *stats§g*_*j*_*|*− *|g*_*i*_
  f. assign *stats§gg*_*IN*_ with *gg*_*IN*_
  g. (g) assign *stats §gg*_*IT*_ with *gg*_*IT*_
  h. Comment - *i*^*th*^ gene influenced
  i. *gg*_*I*_[[*i*]] *<* − list(*ig* = *g*_*i*_, *stats* = *stats*, *PggN* = *PggN*, *PggT* = *PggT*)

